# Uncertainty quantification of parenchymal tracer distribution using random diffusion and convective velocity fields

**DOI:** 10.1101/665109

**Authors:** Matteo Croci, Vegard Vinje, Marie E. Rognes

## Abstract

**Background:** Influx and clearance of substances in the brain parenchyma occur by a combination of diffusion and convection, but the relative importance of thiese mechanisms is unclear. Accurate modeling of tracer distributions in the brain relies on parameters that are partially unknown and with literature values varying up to 7 orders of magnitude. In this work, we rigorously quantified the variability of tracer enhancement in the brain resulting from uncertainty in diffusion and convection model parameters.

**Methods:** In a mesh of a human brain, using the convection-diffusion-reaction equation, we simulated tracer enhancement in the brain parenchyma after intrathecal injection. Several models were tested to assess the uncertainty both in type of diffusion and velocity fields and also the importance of their magnitude. Our results were compared with experimental MRI results of tracer enhancement.

**Results:** In models of pure diffusion, the expected amount of tracer in the gray matter reached peak value after 15 hours, while the white matter does not reach peak within 24 hours with high likelihood. Models of the glymphatic system behave qualitatively similar as the models of pure diffusion with respect to expected time to peak but display less variability. However, the expected time to peak was reduced to 11 hours when an additional directionality was prescribed for the glymphatic circulation. In a model including drainage directly from the brain parenchyma, time to peak occured after 6-8 hours for the gray matter.

**Conclusion:** Even when uncertainties are taken into account, we find that diffusion alone is not sufficient to explain transport of tracer deep into the white matter as seen in experimental data. A glymphatic velocity field may increase transport if a directional structure is included in the glymphatic circulation.

## INTRODUCTION

Over the last decade, there has been a significant renewed interest in the waterscape of the brain; that is, the physiological mechanisms governing cerebrospinal fluid (CSF) and interstitial fluid (ISF) flow in (and around) the brain parenchyma. A number of new theories have emerged including the glymphatic system [37, 39], the intramural periarterial drainage (IPAD) theory [18, 5], and the Bulat-Klarica-Oreskovic hypothesis [53], along with critical evaluations [34, 11, 68]. A great deal of uncertainty and a number of open questions relating to the roles of diffusion, convection and clearance within the brain parenchyma remain.

Exchange between CSF and ISF is hypothesized to occur along small fluid-filled spaces surrounding large penetrating arteries in the brain parenchyma known as paravascular spaces (PVS) [61, 37]. Tracer has been observed to move faster in paravascular spaces in response to increased arterial pulsations, and arterial pulsation has thus been proposed as the main driver of paraarterial flow [30, 38, 48]. After entering the extracellular space (ECS), a bulk flow of ISF from paraarterial to the paravenous spaces has been proposed to occur before re-entry to the the subarachnoid space (SAS) [39]. This concept of CSF/ISF fluid circulation has been named the glymphatic system, with bulk flow as a mechanism for effective waste clearance from the brain parenchyma. Xie et al. [75] showed glymphatic influx to increase in sleeping mice, linking the importance of sleep to clearance of waste products. Sleep was also associated with an increased interstitial space volume fraction, a possible explanation for increased flow through the interstitial space. MRI investigations have also found evidence for glymphatic function in human brains [64, 63].

While several studies demonstrate CSF influx along paraarterial spaces [60, 37, 16, 48], the efflux route is more debated. Carare et al. [18] found evidence of solutes draining from the brain parenchyma along basement membranes of capillaries and arteries, going in the opposite direction of blood flow and possible PVS fluid movement. This flow is however not facilitated by arterial pulsations [23], but by the movement of smooth muscle cells [6]. Bedussi et al. [15] observed tracers move towards the ventricular system, ultimately leaving the brain via the cribriform plate and the nose. A continuous pathway alongside capillaries to the paravenous space has been suggested [31], and capillaries continuously filtrate and absorb water inside the brain parenchyma [53]. In addition, substances may leave the parenchyma crossing the blood-brain barrier, or possibly directly to lymph nodes [35].

In a recent review, Abbott and colleagues [2] concluded that bulk flow within the parenchyma is likely to be restricted to the PVS and possibly white matter tracts, and not present in the neuropil of gray matter. Earlier studies have reported a bulk flow velocity of less than 1 *µ*m/s [51], while recent evidence suggests average net bulk flow of around 20 *µ*m/sec, restricted to the PVS [14, 48]. Nevertheless, since tracer movement in in-vivo studies does not necessarily directly reflect underlying fluid flow [8], the exact velocity field governing ISF flow in the brain remains unknown.

All of the aforementioned in-vivo studies have used tracers or micro-spheres to track the movement of fluid within the intracranial space. Injection of fluid at rates as low as 1 *µ*L/min can cause a significant increase of local intracranial pressure (ICP) [73], which may lead to pressure gradients driving bulk flow. On the other hand, non-invasive methods such as diffusion tensor imaging may serve as a promising tool due to its sensitivity to dispersion and bulk flow. This method has been applied successfully to demonstrate increased diffusivity with vascular pulsation compared to diastole [32]. The diffusion coefficient was found to be anisotropic and highest parallel to PVS, however a value of the bulk fluid velocity could not be reported from these measurements. In addition to both invasive and non-invasive experiments, computational models have been used to assess the possibility and plausibility of bulk flow within the parenchyma. Tracer movement in the extracellular space has been found to be dominated by diffusion [36], a conclusion similar to that of Smith et al. [68] in experimental studies with very low infusion rates.

Even though computational models can distinguish between diffusion and bulk flow, a major challenge remains with regard to the unknown material parameters, boundary conditions and other model configurations needed to accurately predict the movement of ISF in the brain parenchyma. For instance, the permeability of brain tissue used in computational models varies from 10^−10^ to 10^−17^ m^2^ [28, 36]. Because the permeability is directly linked to the Darcy fluid velocity in these models, this parameter choice could result in a difference of 7 orders of magnitude in predicted ISF flow. In addition, CSF dynamics vary between subjects [13] and human CSF production has been reported to increase in the sleeping state [52] which may alter ISF flow. Recently it has been pointed out that there is an overarching need to reduce uncertainty when characterizing the anatomy and fluid dynamics parameters in models considering the glymphatic circulation [66].

Replacing partial differential equation (PDE) parameters subject to uncertainty with spatially correlated random fields is a common modelling choice in the uncertainty quantification (UQ) literature [21, 19, 70] and Monte Carlo methods have been successfully used in biology to quantify how uncertainty in model input propagates to uncertainty in model output. However, these methods have mainly been applied to simulations of the cardiovascular system [57, 17] and, to our knowledge, there has only been one study in which Monte Carlo methods have been used for UQ in brain modelling [33]. To the authors’ knowledge, there has been no previous work on stochastic forward uncertainty quantification for simulations of tracer transport with the brain parenchyma.

With this study, we aim to rigorously quantify how the aforementioned uncertainties in the physiological parameters and in ISF flow affect the spread of a tracer from the SAS into the brain parenchyma. We assume movement of tracer in the brain parenchyma to occur by diffusion and/or convection. To account for uncertainty and variability, we circumvent the lack of precise parameter values by modelling velocity and diffusivity as Matérn stochastic fields. We then set up a PDE model with these stochastic (random) fields as coefficients and quantify the uncertainty in the model prediction via the Monte Carlo (MC) method.

More specifically, we model the contrast MRI study performed by Ringstad et al. [64] assessing glymphatic function in the human brain and derive a baseline convection-diffusion-reaction PDE. The model coefficients are designed to represent different hypotheses on CSF flow and clearance, including diffusion, the glymphatic system and possible capillary absorption, and uncertainty within each hypothesis. A total of five different models were investigated, each with stochastic model coefficients. For each model, we compute the expected values and 99.73% confidence intervals for different functionals of interest of the tracer concentration. The results reported in the study by Ringstad et al. are compared with the range of uncertainty in our model. We find that although the uncertainty associated with diffusion yields great variability in tracer distribution, diffusion alone is not sufficient to explain transport of tracer deep into the white matter as seen in experimental data. A glymphatic velocity field may increase tracer enhancement, but only when adding a directional structure to the glymphatic circulation.

## METHODS

We model the MRI-study of Ringstad et al. [64]. In their experiments, 0.5 mL of 1.0 mmol/mL of the radioactive tracer gadobutrol was injected intrathecally in 15 hydrocephalus patients and eight reference subjects. The localization of the tracer was found with MRI at 4 different time periods, at 1, 3, 4.5, and 24 hours following the injection. After 3 hours, tracer was localized in the lower region of the cranial SAS, and had started to penetrate into the brain parenchyma of the reference subjects. The following day it had spread throughout the brain tissue. Tracer was found to penetrate along large leptomeningeal arteries in all study subjects, and a low proportion of tracer was found at the upper convexities of the brain.

### Gaussian and Matérn fields

Let (Ω, *𝒜*, ℙ) be a probability space, let 𝒟 ⊂ ℝ^3^ be an open domain (representing the brain parenchyma) with coordinates *x* ∈ 𝒟, and let *t* ≥ 0 denote time. A random field *X* = *X*(*x, ω*), *ω* ∈ Ω, *x* ∈ ℝ^*d*^ is a function whose values are random variables for each *x* ∈ ℝ^*d*^. The field is Gaussian if these random variables are all joint Gaussian [3]. A Gaussian field is uniquely determined by providing a mean *µ*(*x*) and a symmetric positive definite covariance function 𝒞 (*x, y*).

A Matérn field is a Gaussian field with covariance of the Matérn class, i.e. of the form

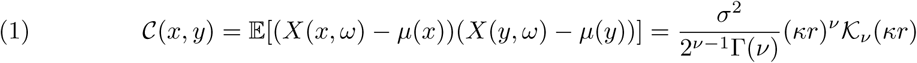

for *x, y* ∈ *𝒟*, where 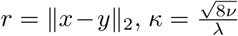, Γ(*x*) is the Euler Gamma function, and *σ*^2^, *ν, λ* > 0 are the variance, smoothness parameter and correlation length of the field respectively and 𝒦_*ν*_ is the modified Bessel function of the second kind. Matérn fields are extensively used in spatial statistics, biology and oil reservoir modelling to represent uncertain or randomly-varying fields [55, 44]. The smoothness parameter *ν* regulates the field’s spatial smoothness: field samples are almost surely continuous and ⌈*ν* ⌉ − 1 times differentiable [3]. For the two cases *ν* = 1*/*2 and *ν* = *∞*, (1) reduces to the exponential and Gaussian covariance kernels, respectively.

The correlation length *λ* roughly represents the distance past which point values of the field are approximately uncorrelated. Informally, this means that in each realization of the Matérn field, there are regions of length proportional to *λ* within which the values of the field are similar.

### Stochastic models for tracer movement in the brain parenchyma

We consider the following partial differential equation with random coefficients to model transport of tracer in the brain parenchyma under uncertainty: find the tracer concentration *c* = *c*(*t, x, ω*) for *x* ∈ *𝒟, ω* ∈ Ω and *t* ≥ 0 such that

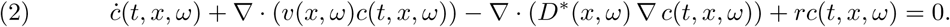

Here, the superimposed dot represents the time derivative, *D*\* is the effective diffusion coefficient of the tracer in the tissue (depending on the tracer free diffusion coefficient and the tissue tortuosity) [51], *v* represents a convective fluid velocity and *r* ≤ 0 is a drainage coefficient potentially representing e.g. capillary absorption [53] or direct outflow to lymph nodes [64]. We assume that the parenchymal domain contains no tracer initially: *c*(0, *x, ω*) = 0.

To investigate and compare different hypotheses for parenchymal ISF flow and tracer transport, we consider 5 stochastic model variations of (2) including two models with stochastic (random) diffusion properties (Model D1 and D2) and three models with stochastic velocity fields (Models V1, V2, and V3). The diffusion-only Models D1 and D2 correspond to negligible ISF bulk flow in the parenchyma and the absence of capillary absorption or other direct outflow pathways. For the velocity models (V1, V2 and V3), we consider a fixed non-random diffusion coefficient in order to isolate the effects of the stochastic velocity fields. A summary of the models are presented in Table 1, while the mathematical modelling aspects are described in further detail in the following sections.

**TABLE 1.** Summary of stochastic model variations with effective diffusion coefficient *D*\*, convective fluid velocity *v*, and drainage coefficient *r* in (2).

| Model | $D^*$ | $v$ | $r$ |
| --- | --- | --- | --- |
| D1 | Random variable | 0 | 0 |
| D2 | Random field | 0 | 0 |
| V1 | Constant | Random influx and outflux field | 0 |
| V2 | Constant | Model V1 + directional velocity field | 0 |
| V3 | Constant | Random influx field | $r < 0$ |

#### Domain and geometry

We define the computational domain 𝒟 as the union of white and gray matter from the generic Colin27 human adult brain atlas FEM mesh [26] version 2 (Figure 1a). This domain includes the cerebellum. The levels of the foramen magnum, the sylvian fissure and the precentral sulcus are well represented by z-coordinates −0.1, 0 and 0.1 m, respectively. The plane z = 0 corresponds approximately to the level of the lateral ventricles.

**FIGURE 1.**
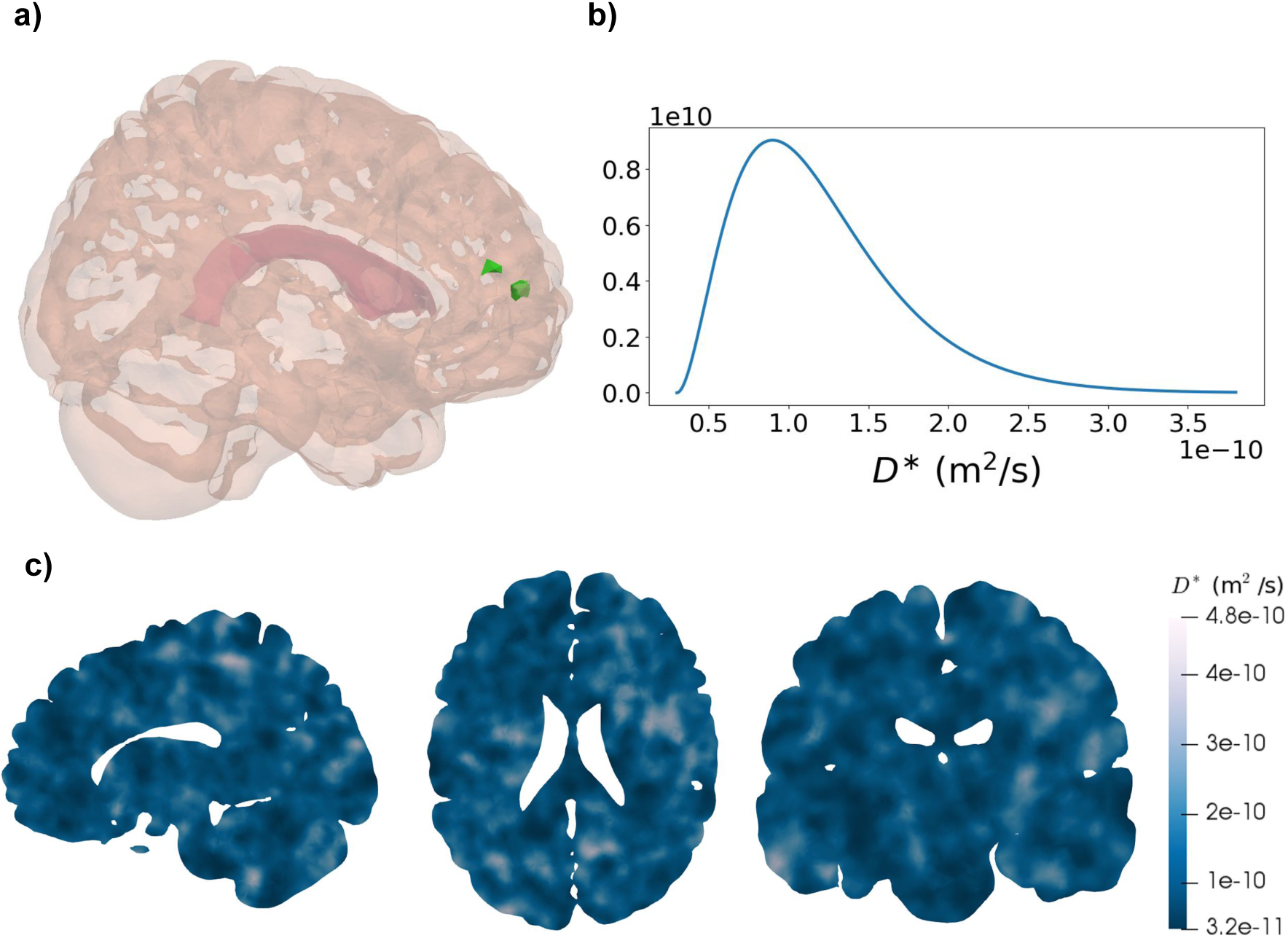
Computational domain and stochastic diffusion coefficient. **a)** The computational domain representing the brain parenchyma including the cerebellum. The interior lateral ventricles are marked (dark pink) in the central region of the domain. Two smaller regions of interest *S*_*g*_ and *S*_*w*_, in the gray and white matter respectively, are marked in green (leftmost region: *S*_*w*_, rightmost region: *S*_*g*_). **b)** Assumed probability distribution of the homogeneous effective diffusion coefficient *D*\* modelled as a random variable and used in Model D1. The expected value *E*[*D*\*] is 1.2 ×10^−10^ m^2^/s. **c)** Sample of the heterogeneous effective diffusion coefficient (sagittal, axial and coronal slices ordered from left to right) modelled as a random field and used in Model D2.

#### Boundary conditions modelling tracer movement in the SAS

Let *∂D* be the boundary of *𝒟* and let *∂𝒟* = *∂𝒟*_*S*_ ∪ *∂𝒟*_*V*_, with *∂𝒟*_*S*_ representing the interface between the brain parenchyma and the subarachnoid space (SAS), and *∂𝒟*_*V*_ representing the interface between the brain parenchyma and cerebral ventricles, respectively. We consider the following boundary conditions for (2):

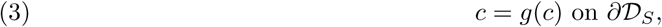

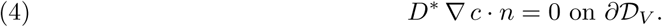

In particular, we assume that a tracer concentration is given at the SAS interface (3) and no ventricular outflux (4). The dependence of *g* on *c* in (3) is detailed below.

The boundary condition (3) models the movement of tracer starting from the lower cranial SAS and traveling upward in the CSF surrounding the brain as observed in the study by Ringstad et al [64]. In particular, we let

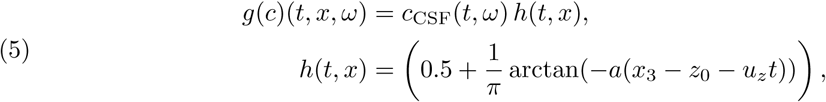

for *x* = (*x*_1_, *x*_2_, *x*_3_). Here, at time *t, c*_CSF_(*t*) is the average tracer concentration in the SAS, while *h*(*t, x*) represents its spatial distribution.

The expression for *h* is based on the following considerations. We assume that the diffusive and/or convective movement of tracer from the spinal to the cranial SAS over time is known, and we thus model *h*(*t, x*) as a smooth step function upwards (in the *x*_3_- or *z*-direction). In (5), *u*_*z*_ represents the speed of tracer movement upwards in the SAS, and *a* reflects the gradient of tracer concentration from the lower to the upper cranial SAS. Finally, we assume that at time *t* = 0, the tracer has spread up to a relative distance of *z*_0_ from the lateral ventricles. This specific expression for *h*(*t, x*) and the values of parameters *a, z*_0_ and *u*_*z*_ are based on the spread of tracer seen in the MR-images in the study by Ringstad et al. [64]. In particular, we use *a* = 20 m^−1^, *u*_*z*_ = 1.5 × 10^−5^ m/sec and *z*_0_ = −0.2 m. These parameters were chosen to match time to peak in three different regions in the CSF space in reference individuals [64].

To derive an expression for *c*_CSF_ in (5), we consider the conservation of tracer mass. We model the spread of *n*_0_ = 0.5 mmol tracer in the CSF, assuming a volume of *V*_CSF_ = 140 mL CSF in the human SAS and ventricles [74]. The average concentration in the SAS right after injection is thus *c*_CSF_(0) = 0.5 mmol/140 mL = 3.57 mol/m^3^. At any given time, we assume that the total amount of tracer in the brain and in the SAS plus or minus the tracer absorbed or produced stays constant in time, and is equal to the initial amount *n*_0_ = 0.5 mmol (almost surely):

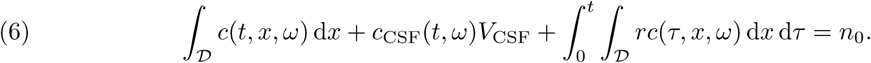

Solving for *c*_CSF_, we thus obtain

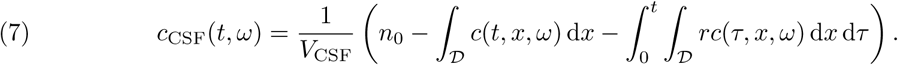

#### Quantities of interest

To evaluate the speed and characteristics of tracer movement into and in the brain parenchyma, we consider a set of functionals describing different output quantities of interest. To quantify the overall spread of tracer in the gray and white matter, we consider the (integrated) amount of tracer in the gray matter *Q*_*g*_ and in the white matter *Q*_*w*_ at time points *τ* :

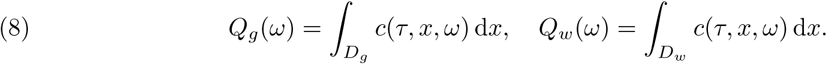

We pay particular attention to the times *τ* ∈ {3, 5, 8, 24}. To further differentiate, we also defined two localized functionals at each time *τ* : the average tracer concentration *q*_*g*_ in a small subregion of the gray matter *S*_*g*_ and analogously *q*_*w*_ for a small subregion of the white matter *q*_*w*_:

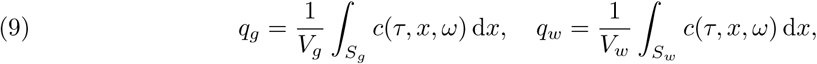

where *V*_*g*_ and *V*_*w*_ is the volume of the gray and white matter subregions, respectively. The size and relative location of the subregions *S*_*g*_ and *S*_*w*_ within the computational domain are illustrated in Figure 1a. To further quantify the speed of propagation, we define the white matter activation time *F*_*w*_:

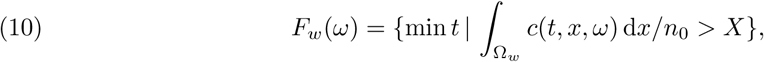

where *n*_0_ is the total amount of tracer injected into the SAS (0.5 mmol) and *X* is a given percentage. We here chose *X* = 10%. Finally, we also define the analogous regional (white matter) activation time

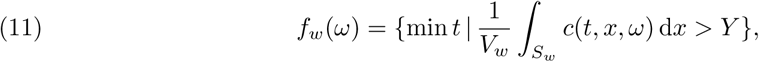

where *Y* = 10^−3^ mol/m^3^

For plotting the boundary tracer concentration over time, we define three axial planes along the z-axis (*z* = −0.1, 0, 0.1 m) to represent the level of the foramen magnum, sylvian fissure and precentral sulcus, respectively.

### Stochastic diffusion modelling

The parenchymal effective diffusion coefficient of a solute, such as e.g. gadobutrol, is heterogeneous [72] (varies in space) and individual-specific (varies from individual to individual). To investigate the effect of uncertainty in the diffusion coefficient, we consider two approaches: first, to model the diffusion coefficient as a random variable and second, to model the diffusion coefficient as a random field, thus allowing for tissue heterogeneity. Both approaches are described in further detail below.

#### Effective diffusion coefficient modelled as a random variable

First, we consider the simplifying but common assumption that the effective diffusion coefficient is spatially homogeneous: *D*\*(*ω*) ∈ℝ. We account for the uncertainty in its value by modelling it as a random variable:

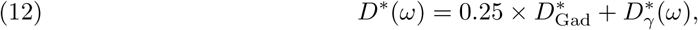

where 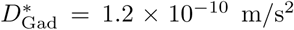 is a fixed parenchymal gadobutrol diffusivity [63] and where 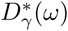 has a Gamma distribution with shape *k* = 3 and scale 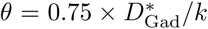. The choice of shape and scaling parameters ensures that (i) the diffusion coefficient is positive, (ii) its expected value matches reported values of parenchymal gadobutrol diffusivity [63], and (iii) its variability allows for values up to 2–3 times larger or smaller than the average with low probability. The last modelling choice reflects diffusivity values in the range 1-10 ×10^−10^ m/s^2^ in agreement with previous reports [51]. The probability distribution of *D*\* is shown in Figure 1b-c.

#### Effective diffusion coefficient modelled as a random field

In order to represent spatial heterogeneity in the diffusion coefficient, we next model *D*\* as a continuous random field. Again, we set

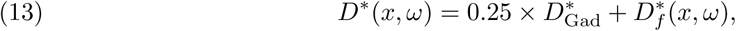

where 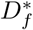 now is a random field such that for each fixed 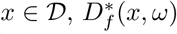is a gamma-distributed random variable with the same parameters as *D*\*(*ω*) in (12). To enforce continuity and to easily sample the random field from its distribution, we draw samples of 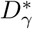 by first sampling a Matérn field *X*(*x, ω*) and then transforming it into a gamma random field by using a copula [50]. This consists in setting 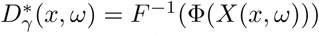, where *F* ^−1^ is the inverse cumulative density function (CDF) of the target (gamma) distribution, Φ is the CDF of the standard normal distribution and *X*(*x, ω*) is a standard (zero mean, unit variance) Matérn field with smoothness parameter *ν* = 2.5 and correlation length *λ* = 0.01 m, cf. (1). Note that spatial changes in the diffusivity occurs at a length scale corresponding to the correlation length, here 0.01 m.

### Stochastic velocity modelling

In what follows we introduce three different models for the velocity field, each representing a different hypothesis regarding intraparenchymal ISF/CSF movement. We emphasize that each model represent a homogenized velocity field averaged over physiological structures.

#### Glymphatic velocity model: arterial influx and venous efflux

To define a stochastic homogenized velocity model representing the glymphatic pathway, we assume that ISF follows separate inflow and outflow routes: entering the brain along paraarterial spaces and exiting along paravenous spaces [39]. We further suggest that

1. Substantial changes within the velocity field happen after a distance proportional to the mean distance between arterioles and venules.
2. The blood vessel structure is random and independent from the position within the parenchyma in the sense that the presence of paraarterial or paravenous spaces are equally likely at any point in space. Mathematically, this assumption requires the expected value of each of the velocity components to be zero.
3. The velocity field varies continuously in space and is divergence-free (∇·*v* = 0), i.e. no CSF/ISF leaves the system e.g. through the bloodstream.
4. We set the expected velocity magnitude 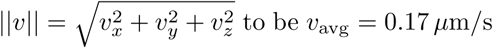 and we allow for up to 2-3 times larger and up to 10 times smaller values with low probability [51].

Although ISF/CSF velocities in paravascular regions may be higher [48] that what we propose, the velocity field here models an averaged bulk flow over a larger area (comprised of e.g. PVS and adjacent tissue). Bulk flow velocities in rats have been reported to be in the range of approximately 0.1-0.24 *µ*m/s [1, 51].

To address these stipulations, we define the stochastic glymphatic circulation velocity field

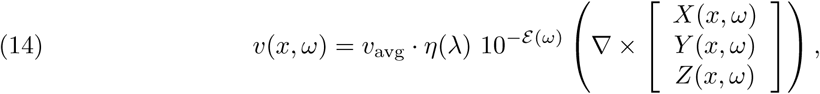

where 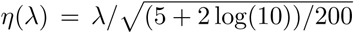 is a scaling constant chosen such that the magnitude of *v* satisfies 𝔼[‖*v*‖ ^2^]^1*/*2^ = *v*_avg_ (we omit the mathematical derivation of this constant), *ε* (*ω*) is an exponentially distributed random variable with mean 0.2 and *X*(*x, ω*), *Y* (*x, ω*) and *Z*(*x, ω*) are standard independent identically distributed (i.i.d) Matérn fields with *ν* = 2.5 and correlation length *λ* = 1020 *µ*m. A sample of the glymphatic circulation velocity field together with the velocity magnitude distribution is shown in Figure 2a-b.

**FIGURE 2.**
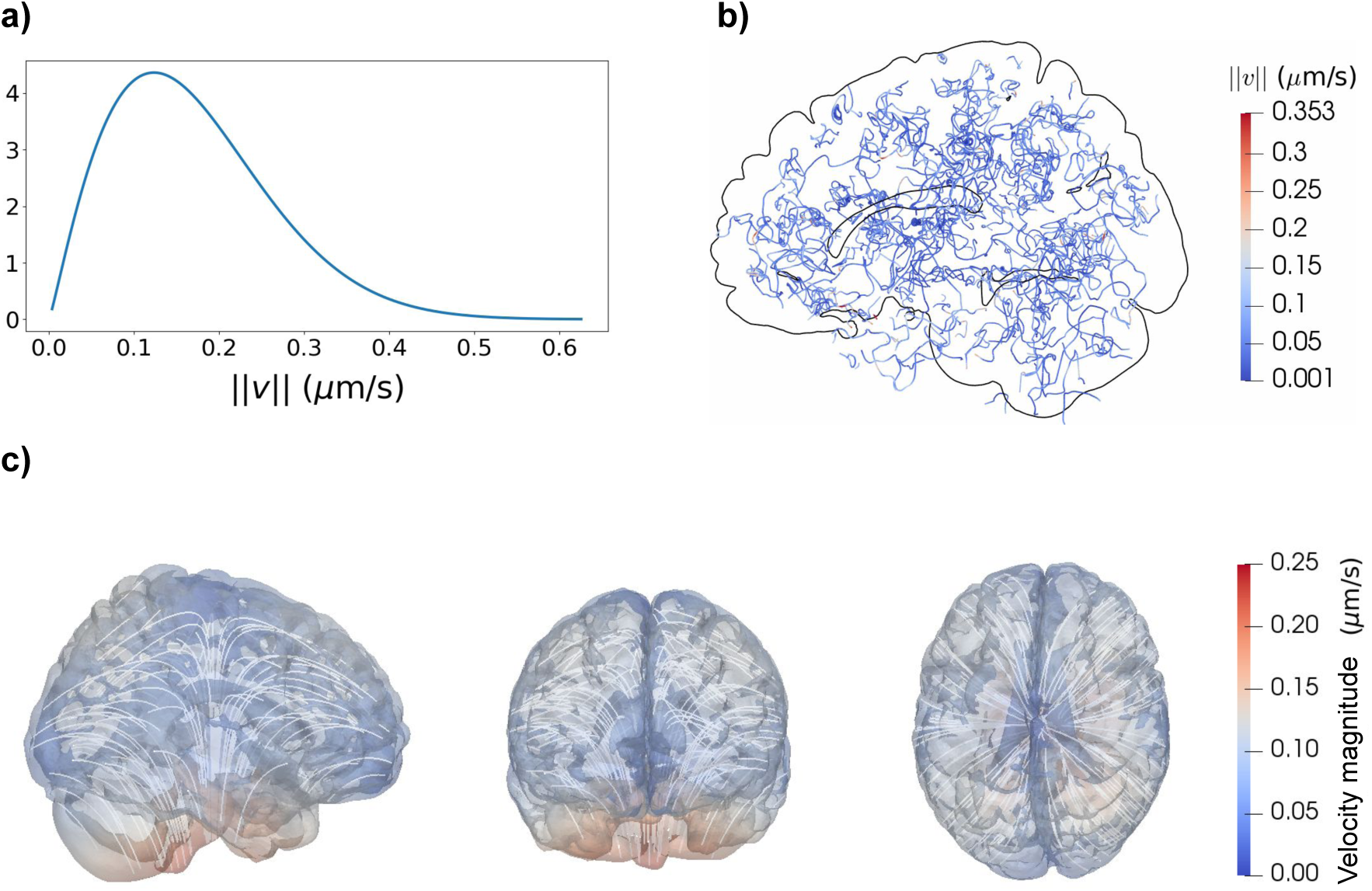
Stochastic aspects of the glymphatic circulation velocity fields (Models V1 and V2). **a)** Probability density of the glymphatic circulation velocity magnitude ‖*v*‖ cf. (14). **b)** Streamlines of a corresponding velocity field sample. **c)** Velocity magnitude and streamlines for the directional velocity field *v*_dir_ as given by (15). The flow field is assumed to follow cardiovascular pulses upwards along the brain stem. After entering the deeper parts of the brain, the bulk flow spreads out at reduced velocity. From left to right: sagittal, coronal and transverse view.

The factor 10^*-ε* (*ω*)^ is an ad-hoc random term to enforce the variability requirement defined by point 4) above. The use of Matérn fields enforces spatial variability in a continuous manner and taking the curl operator (∇×) ensures that the resulting velocity is divergence-free, hence addressing point 3). It can be proven (although we omit the details here) that the field within the brackets in (14) is still Gaussian, has zero mean (hence satisfies 2)) and has the same correlation length as the original Matérn fields, albeit it presents a slightly different covariance structure.

The choice of correlation length was guided by the following considerations. The mean distance between arterioles and venules was reported to be 280 *µ*m in rhesus monkeys [4], although the value 250 *µ*m has been used as a representative distance in humans in recent modeling papers [40, 59]. We estimated the mean distance in humans by considering differences in brain and artery size between monkey and human (Table 2). We find a factor close to 2 between CCA and arteriole diameter, while a similar ratio was found for the cube root of the brain mass. Thus, the correlation length should be greater than 250 − 560*µ* m to address point 1) above. Combining these physiological considerations with the corresponding requirements on the numerical resolution, we let *λ* = 1020 *µ*m.

**TABLE 2.** Brain-related parameters of three species. *: Estimated values. *d*_CCA_: diameter of the common carotid artery, *d*_A_: arteriole diameter Δ_AV_: distance between arteriole and venule.

| Species | Brain mass [g] | $d_{CCA}$ [mm] | $d_A$ [$\mu\text{m}$] | $\Delta_{AV}$ [$\mu\text{m}$] |
| --- | --- | --- | --- | --- |
| Mouse | 0.3 [65] | 0.47 [43] | 25 [38] | 40* |
| Monkey | 88 [65] | 3.5 [76] | 35.5 [4] | 280 [4] |
| Human | 1350 [65] | 6.3 [42] | 40-250 [10] | 1020* |

#### Glymphatic velocity model with additional directional velocity field

Above we assumed that the blood vessel distribution was independent of the spatial position within the parenchyma and that bulk flow from arterial to venous PVS occurs on a small length scale proportional to the mean distance between arterioles and venules. However, transport of tracer might also happen on a larger length scale along larger vascular structures present in given physical regions such as e.g. circle of Willis). AS CSF is hypothesized to enter the brain along penetrating arteries, the direction of cardiac pulse propagation may induce a directionality of the glymphatic circulation as well. The cardiac pulse follows the vessel paths of larger arteries entering the brain from below, and from there spreads out almost uniformly [41, 58]. The pulses also seem to traverse deep gray matter structures on the way up towards the ventricles.

To model such behavior, we introduce a directional velocity field *v*_dir_, with characteristics qualitatively similar to what is described in the literature [41, 58]:,

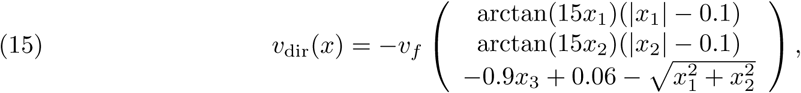

where *v*_*f*_ = 2 ×10^−6^ m/s. For a plot of *v*_dir_, see Figure 2c. The velocity field *v*_dir_ induces a net flow out of the parenchyma at the very low rate of 0.007 mL/min. We superimpose this deterministic directional velocity field by the stochastic glymphatic circulation velocity field to define the stochastic glymphatic directional velocity field:

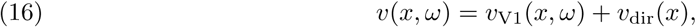

where *v*_V1_ is given by (14). This velocity model thus takes into account both the “randomness” of small arteries, but also the “deterministic” presence of large arteries and possibly other structures of blood flow propagation [41, 58].

#### Capillary filtration model V3: arterial inflow with a homogeneous sink throughout the brain

Several independent studies demonstrate that CSF may enter the brain parenchyma along spaces surrounding penetrating arteries [48, 39, 5, 14]. However, the glymphatic efflux concept of a bulk flow of CSF through the ECS and recirculation into the SAS through paravenous spaces has been severely questioned [34, 36, 5, 67]. As a variation, we here therefore also consider a stochastic velocity model representing paraarterial influx without a direct return route to the CSF. Instead, we assume that ISF/CSF is drained inside the brain parenchyma along some alternative efflux pathway. This pathway may include the capillaries or separate spaces along the PVS directly into cervical lymph nodes.

In light of this, we consider the following alternative velocity assumptions. (1) There is a net flow of CSF into the brain and (2) ISF is cleared within the parenchyma via some, here unspecified, route. For instance, it has been proposed that production and absorption is present all over the CSF system and that capillaries and ISF continuously exchanges water molecules [54]. However, drainage of large molecules through this route is unlikely as capillaries and the basement membranes are connected through tight junctions [34]. It has also been reported that lymph vessels may be capable of also draining larger molecules from brain tissue into deep cervical lymph nodes, possibly through paravenous spaces [9]. In addition, other outflow routes may exist, including degradation clearance and meningeal lymphatic vessel clearance [69].

To address these assumptions, we define a stochastic arterial inflow velocity field as a radially symmetric field pointing inwards from the SAS interface to the brain region around the lateral ventricle. This central region is modelled in what follows as a sphere of radius *R* = 8 cm and center given by *x*_*c*_ in the lateral ventricles. Mathematical experimentation lead to the following *ansatz* for such velocity:

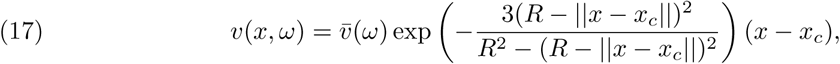

where 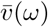 is a gamma random variable chosen such that the probability distribution of the velocity magnitude is comparable to that of the glymphatic circulation velocity defined by (14). The shape parameter *k* = 2 and the scale parameter is set such that again E[‖*v*‖^2^]^1*/*2^ = *v*_avg_. Note that in this case, the expected value of the velocity components are non-zero. To satisfy (2), we model the drainage of tracer by setting *r* = −1 ×10^−5^ s^−1^, which typically results in 40% drainage of the injected tracer over 48 hours.

### Random field sampling and uncertainty analysis

We considered six output functionals of interest: the amounts of tracer in gray and white matter at given times (8), the average tracer concentrations in subregions of gray and white matter (9), the white matter activation time (10), and the white regional activation time (11). Each functional *Q* = *Q*(*ω*) depends on the random parameter *ω* via *c*(·, ·, *ω*) as defined by (2). To sample the functional from its distribution, we first compute a sample of each of the random coefficients in (2) from their distribution, second, solve (2) with the given coefficient sample, and third, evaluate the functional with the computed solution. For sampling the random diffusion and velocity coefficient fields, we adopted a white noise sampling technique using an auxiliary extended domain [22]. We used the standard Monte Carlo approximation to estimate the expected functional value 𝔼[*Q*]

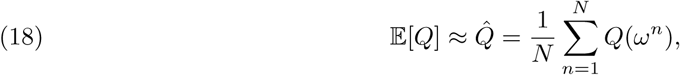

where *N* is the number of Monte Carlo samples. The statistical error introduced by approximating 𝔼[*Q*] with 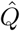 decreases with *O*(*N* ^−1*/*2^). We let *N* = 3200 to ensure an that 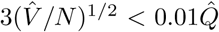, where 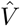 is the sample variance of 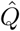.

### Numerical methods and implementation

The diffusion-convection equation (2) was solved numerically using a finite element method with continuous piecewise linear finite elements in space, and a second-order (implicit midpoint) finite difference discretization time with time step Δ*t* = 15 min, combined with mass lumping [71]. The finite element mesh 𝒯_*h*_ was an adaptively refined version of the gray and white matter of the Colin27 human adult brain atlas mesh [26] version 2 with 1 875 249 vertices and 9 742 384 cells. An outer box of dimensions 0.16 × 0.21 × 0.17 (m^3^) with mesh size 0.0023 m was used for the sampling of the Gaussian fields.

For the models with non-zero velocity (Models V1, V2, V3), (2) was typically mildly convection-dominated with an upper estimate of the Péclet number of

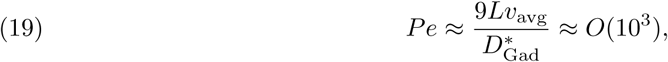

where *L* ≈ 0.084 m is half the diameter of the computational domain, *v*_avg_ = 0.17*µ*m/s, and 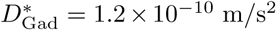. The boundary condition (7) was discretized explicitly in time using the trapezoidal rule; i.e. we let for each n = 0; 1 …:

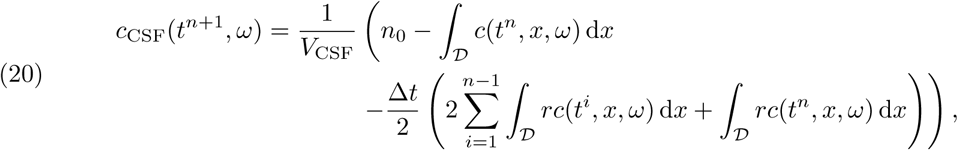

where *t*^*n*^= *n*Δ*t* and the term in the inner bracket results from the numerical integration of the term 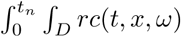.

The numerical solver was verified using a convergence test comparing different mesh refinements, time steps, and stabilization techniques, including SUPG [24], for a set of deterministic worst-case models (with large velocities and small diffusion coefficients) (Additional file 1).

The numerical solver was implemented in Python using the FEniCS finite element software [7] and previously verified in-house parallel Monte Carlo routines [22]. The extended box mesh was created using the Gmsh software [27]. The linear system was solved using the PETSc [12] implementation of the GMRES algorithm preconditioned with the BoomerAMG algebraic multigrid algorithm from Hypre [25]. We used Matplotlib (version 2.1.1) and Paraview (version 5.4.1) for visualization.

## RESULTS

### Non-random diffusion as a baseline for parenchymal solute transport

To establish a baseline for parenchymal solute transport, we first simulated the evolution of a tracer spreading in the SAS and in the parenchyma via diffusion only, using a constant (i.e. non-random) effective diffusion coefficient for gadobutrol (*D*\* = 1.2 × 10^−10^ m^2^/s). The resulting parenchymal tracer spread over 24 hours is shown in Figure 4. The tracer concentration increases first in inferior regions and in the gray matter. Tracer does not penetrate deep into white matter regions within this time frame. In the sagittal plane (top), tracer enhancement is more prominent than in the other two plane as the sagittal plane shown is close to the CSF-filled longitudinal fissure.

**FIGURE 3.**
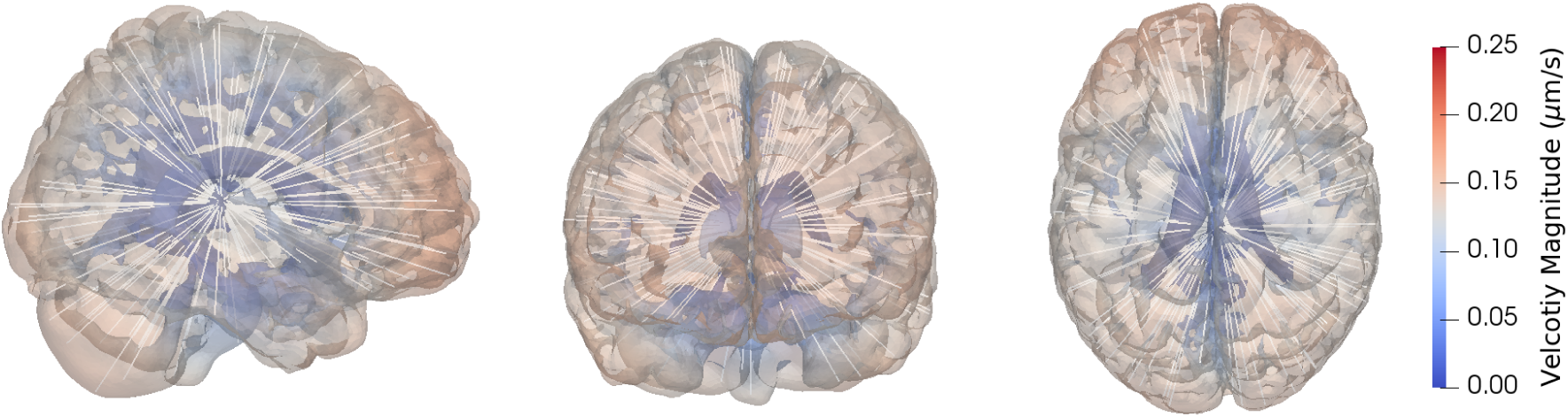
Sample Model V3 velocity field. Velocity magnitude and stream-lines for the velocity field as given by (17). Flow is assumed to occur from the cortex towards the ventricles with reduced velocity along the way due to clearance. From left to right: sagittal, coronal and transverse view.

**FIGURE 4.**
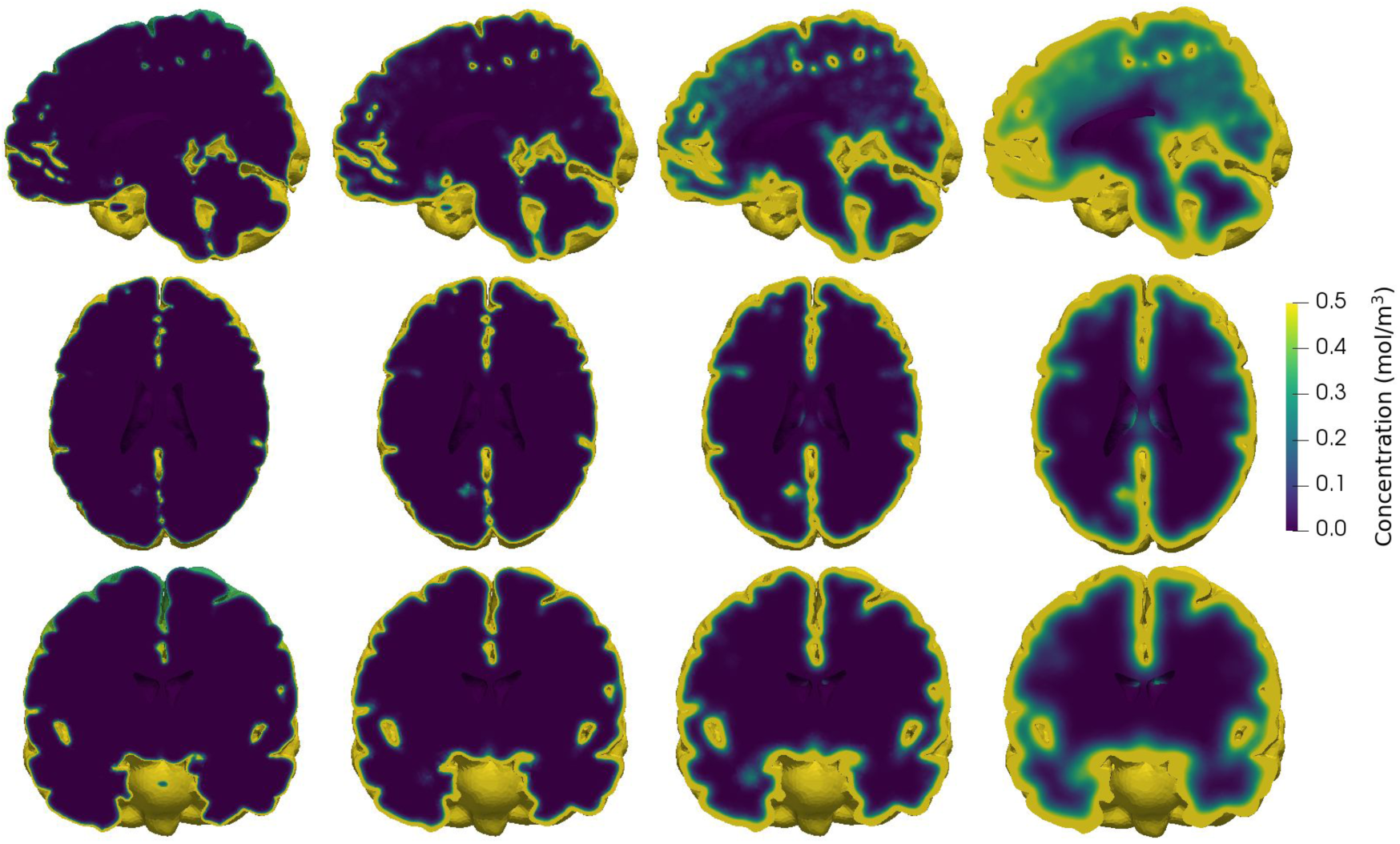
Baseline tracer evolution. Parenchymal tracer concentration after (from left to right) 1, 3, 8 and 24 hours of diffusion in (from top to bottom) sagittal, transverse and coronal planes. Initially, most of the tracer is found in inferior regions. At 24 hours, tracer has penetrated substantially into the gray matter, but not into the deep, central regions.

Figure 5a shows the boundary tracer concentration (concentration in the SAS) over time at the levels of the foramen magnum (*z* = −0.1 m), sylvian fissure (*z* = 0 m) and precentral sulcus (*z* = 0.1 m). During the first few hours, boundary tracer concentration at the level of the foramen magnum increases rapidly, and peaks at 3 hours reaching approximately 2.0 mol/m^3^. Boundary tracer concentrations close to the sylvian fissure and precentral sulcus are lower, and the time to reach peak concentrations is longer. For the sylvian fissure, peak concentration in the CSF is 1.4 mol/m^3^, at 5 hours, while the precentral sulcus concentration reaches 1.1 mol/m^3^ at 7 hours. We note that as the boundary condition depends on the parenchymal tracer concentration itself (cf. (7)), the boundary tracer concentration will differ slightly in subsequent simulation setups.

**FIGURE 5.**
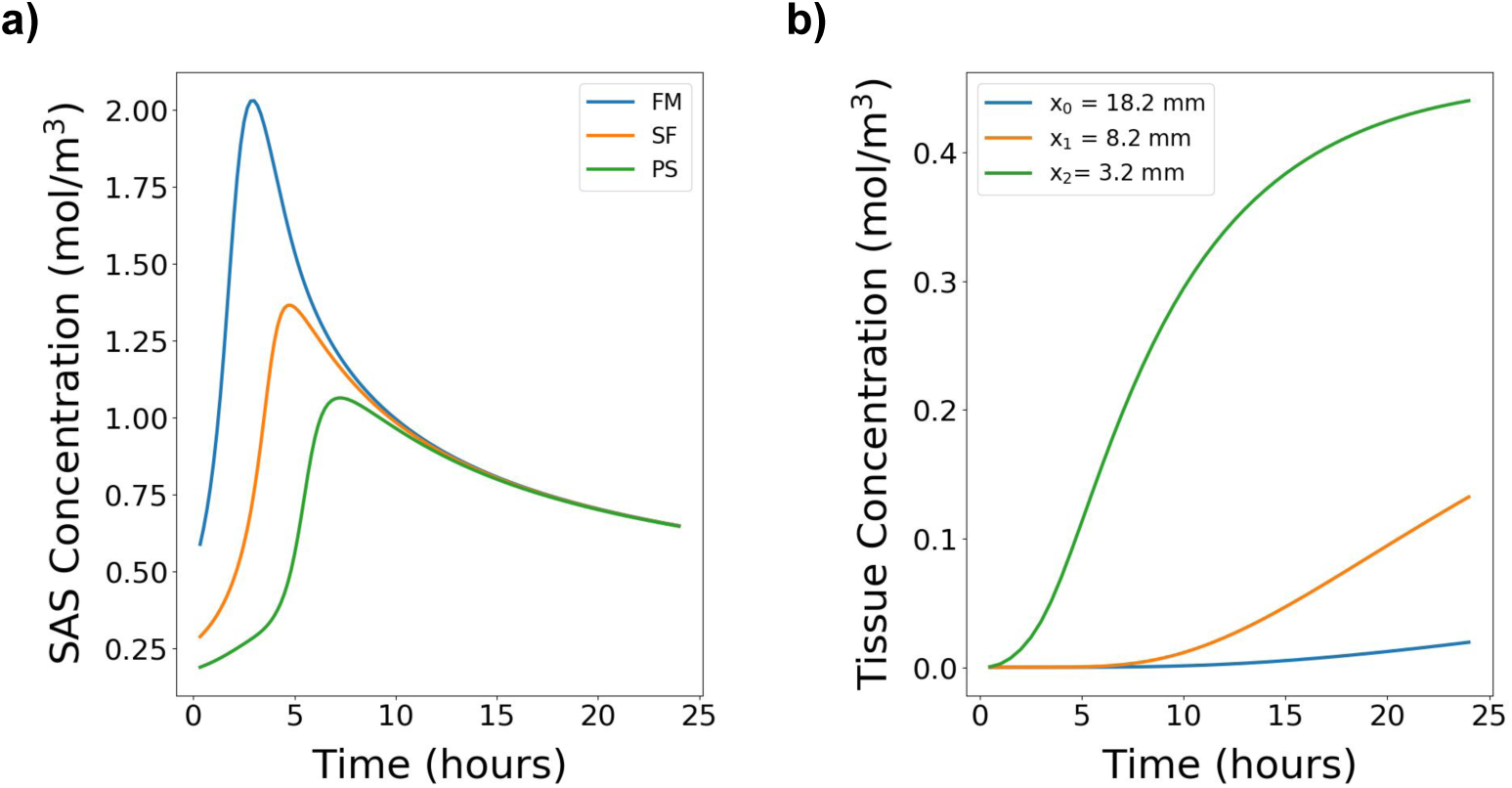
Tracer concentrations. (a) Tracer concentration in the SAS (*c*_CSF_) used as boundary conditions at the brain surface at the level of the foramen magnum (FM), sylvian fissure (SF) and the precentral sulcus (PS). At the lower level of the SAS, tracer concentration peaks at around 3 hours, while at the upper levels, peak concentration occurs later. Following peak values, the concentration in the SAS decreases as tracer enters the parenchyma. The SAS concentration is modeled by (5) (b) Tracer concentration over time in three different points at a given distance from the brain surface. The points were chosen along a line directly from the cortex towards the ventricles at the level of the sylvian fissure.

In Figure 5b, concentration profiles are shown for three interior points at different distances from the brain surface. The points were chosen along a line from the brain surface towards the ventricles at the height of the sylvian fissure (z = 0). The tracer concentration at these points stays low for the first few hours before steadily increasing. For the point closest to the SAS (*x*_2_), the concentration rises faster than for the other two points, and is almost equal to the SAS concentration at 24 hours (0.4 vs 0.5 mol/m^3^). In the middle point (*x*_1_), tracer concentration starts increasing after 6-7 hours and reaches approximately 0.15 mol/m^3^ after 24 hours. For the most interior point (*x*_0_), tracer concentration starts and stays low throughout the 24 hour time span. At 24 hours, the tracer concentration in all three points is still increasing.

### Quantifying the effect of uncertainty in effective diffusion magnitude

We first aimed to quantify the effect of uncertainty in the magnitude of the effective diffusion coefficient on the time evolution of tracer in the gray and white matter. In particular, we computed the tracer concentration, together with auxiliary output quantities, evolving via diffusion only with a gamma-distributed random variable diffusion coefficient (Model D1).

The amount of tracer found in the gray and white matter differ both in magnitude and variation (Figure 6a-c). The expected amount of tracer in the gray matter increases rapidly, and doubles from 1 to 2 hours (0.065 to 0.13 mmol), and again from 2 to 4 hours (0.13 mmol to 0.25 mmol). The gray matter reaches a peak after approximately 15 hours, while the white matter did not reach steady steady within 24 hours. There is substantial variation in the amount of tracer in gray matter throughout the 24 hour time span. The variation is at its largest between 2 and 8 hours where the length of the 99.73%-intervals range from 0.064 mmol to 0.11 mmol corresponding to 13-22% of the total tracer injection of 0.5 mmol. Ultimately, the amount of tracer will reach a steady-state solution, constant in space and time, independently of the diffusion coefficient. Therefore, after a certain point in time, variation decreases as all solutions converge towards the same steady state. The changes in variation of tracer found in the gray matter over the 24 hours are also illustrated by the change in the estimated probability density function (PDF) of the total amount of tracer at a given time (Figure 6c). After 3 and 5 hours (blue and orange curve) the PDFs are symmetric, and with more spread for the later time point. As time evolves, the PDFs become more left skewed (green and red curve), as in almost all cases, the concentration approaches but never surpasses the steady state value.

**FIGURE 6.**
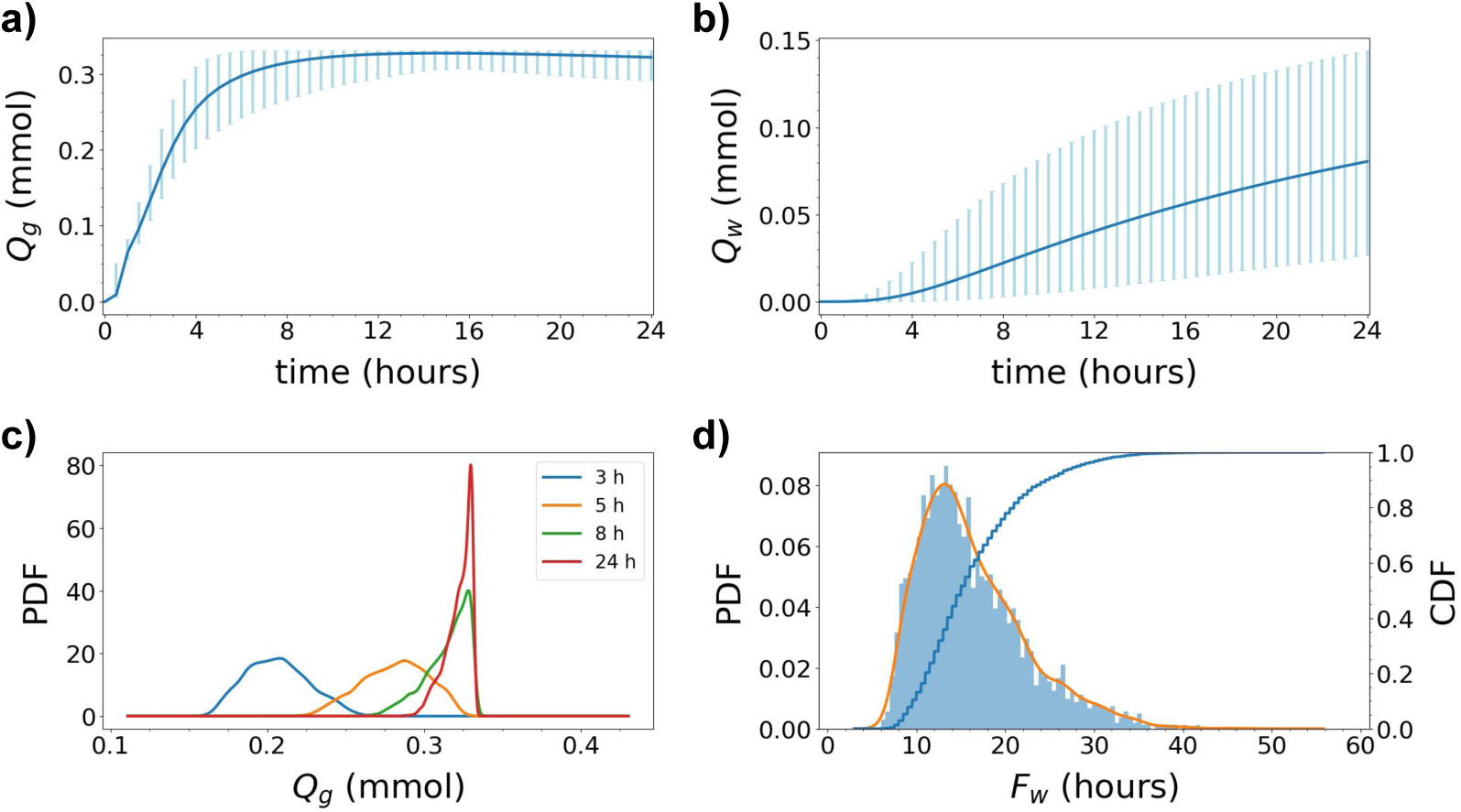
Uncertainty quantification for Model D1. The integrated amount of tracer in the (a) gray matter *Q*_*g*_ and (b) white matter *Q*_*w*_ over time; *Q*_*g*_ and *Q*_*w*_ as defined by (8). The blue curves show the expected value. The light blue vertical bars indicate the variability: 99.73% of the samples fall within the plotted range (with 0.135% of the samples above and 0.135% below). (c) The probability density functions (PDFs) corresponding to *Q*_*g*_ at 3, 5, 8 and 24 hours after tracer injection. (d) Histogram of white matter activation time *F*_*w*_ as defined by (10) (bars), corresponding estimated PDF (orange curve), and corresponding cumulative density function (CDF). Uncertainty in the magnitude of the effective diffusion coefficients substantially impact the amount of tracer found in the gray and white matter and the white matter activation time.

The amount of tracer in the white matter changes slowly for the first two hours, before starting to increase after 3-4 hours (Figure 6b). After 4 hours, the expected amount of tracer in the white matter is only 0.0048 mmol, increasing to 0.022 mmol after 8 hours, and 0.056 mmol after 16 hours. The variation is substantial and increasing with time: the length of the 99.73%-interval is 0.022 mmol at 4 hours, 0.065 mmol at 8 hours and 0.10 at 16 hours. At 24 hours, the uncertainty in diffusion coefficient may explain a factor of approximately 5 in deviation from the lowest (0.027 mmol) to the highest (0.14 mmol) predicted amount of tracer in the white matter.

The estimated PDF and cumulative density function (CDF) for the white matter activation time (i.e. time for 10% of tracer to reach the white matter) is shown in Figure 6d. We observe that the most likely white matter activation time is approximately 14 hours. The white matter activation time is less (than 10%) likely to be less than 9.5 hours, but (more than 90%) likely to be less than 24.5 hours. The activation time may exceed 24 hours, but is highly unlikely to go beyond 40 hours (CDF ¿ 0.998). The white matter activation threshold was reached in all samples within the simulation time span.

### Quantifying the effect of uncertainty in diffusion heterogeneity

Brain tissue is hetero-geneous [72], varies from individual to individual, and is clearly not accurately represented by a single diffusion constant. To further investigate the effect of uncertainty in the diffusion coefficient and in particular to study the effect of spatial heterogeneity, we modelled the diffusion coefficient as a spatially-varying random field (Model D2).

The amounts of tracer found in gray and white matter for Model D2 are nearly identical to those resulting from Model D1 in terms of expected value (data shown later cf. Figure 9), but with substantially less variability. The length of the 99.73% confidence interval for amount of tracer in gray matter (*Q*_*g*_) is less than 0.0071 mmol for all times after the first half hour, corresponding to a relative variability (compared to the expected value) of between 2.2 and 10.9% throughout the 24 hour time span. For white matter, the length of the 99.73% confidence interval is increasing with time, with the relative variability at 24 hours at 7.9%.

**FIGURE 7.**
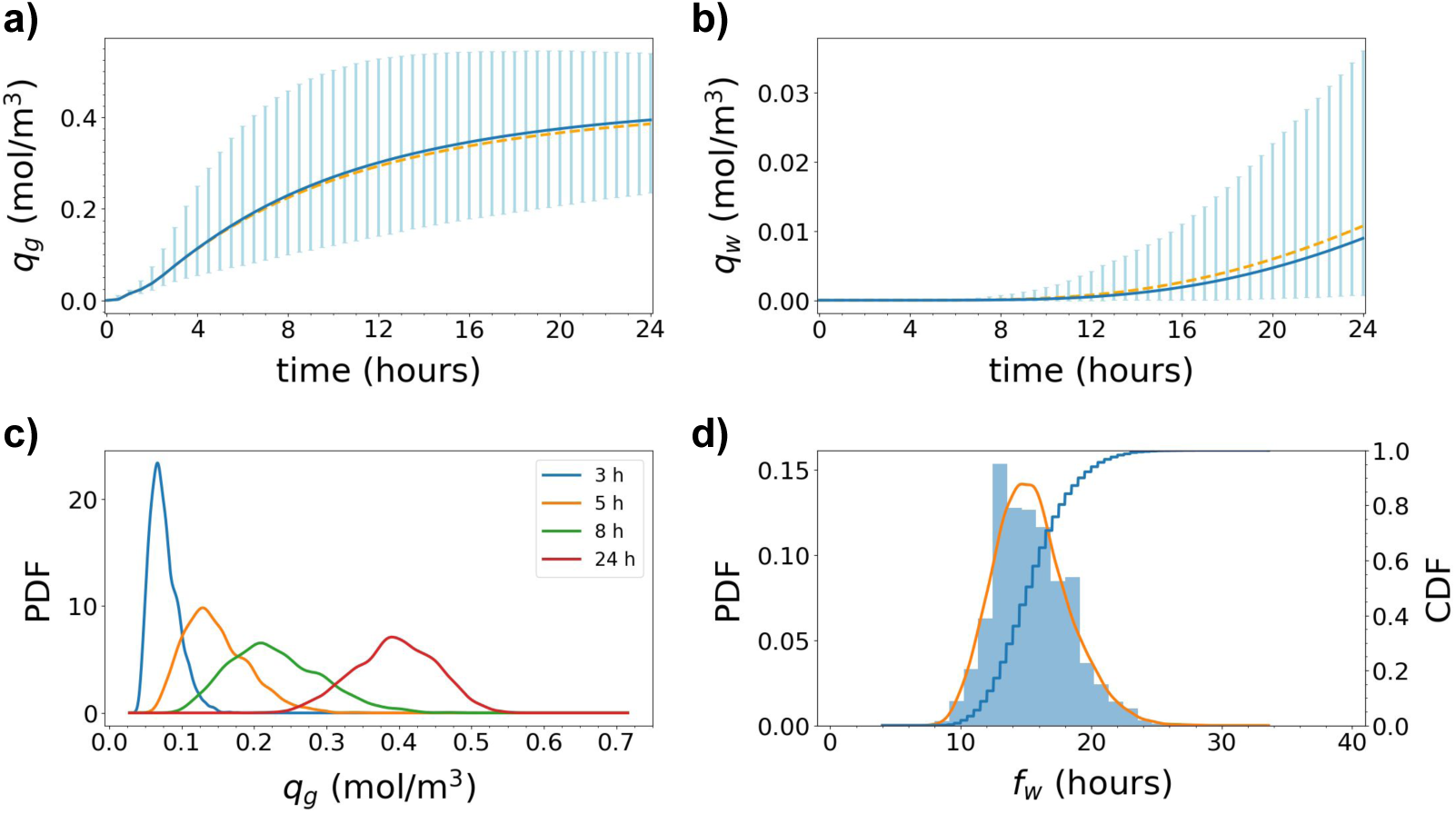
Uncertainty quantification for Model D2. The average tracer concentration in a subregion of (a) gray matter *q*_*g*_ and (b) white matter *q*_*w*_ as defined by (9). The blue curves show the expected value. The light blue vertical bars indicate the variability: 99.73% of the samples fall within the plotted range (with 0.135% of the samples above and 0.135% below). The dashed orange lines in (a) and (b) indicate the analogous expected value curve resulting from Model D1 (constant diffusion only), for comparison. (c) The probability density functions (PDFs) corresponding to *q*_*g*_ at 3, 5, 8 and 24 hours after tracer injection. (d) Histogram of white subregion activation time *f*_*w*_ as defined by (11) (bars), corresponding estimated PDF (orange curve), and corresponding cumulative density function (CDF). Uncertainty in the heterogeneity of the diffusion coefficient leads to a wide range of likely average tracer concentrations in the white matter throughout the time span.

**FIGURE 8.**
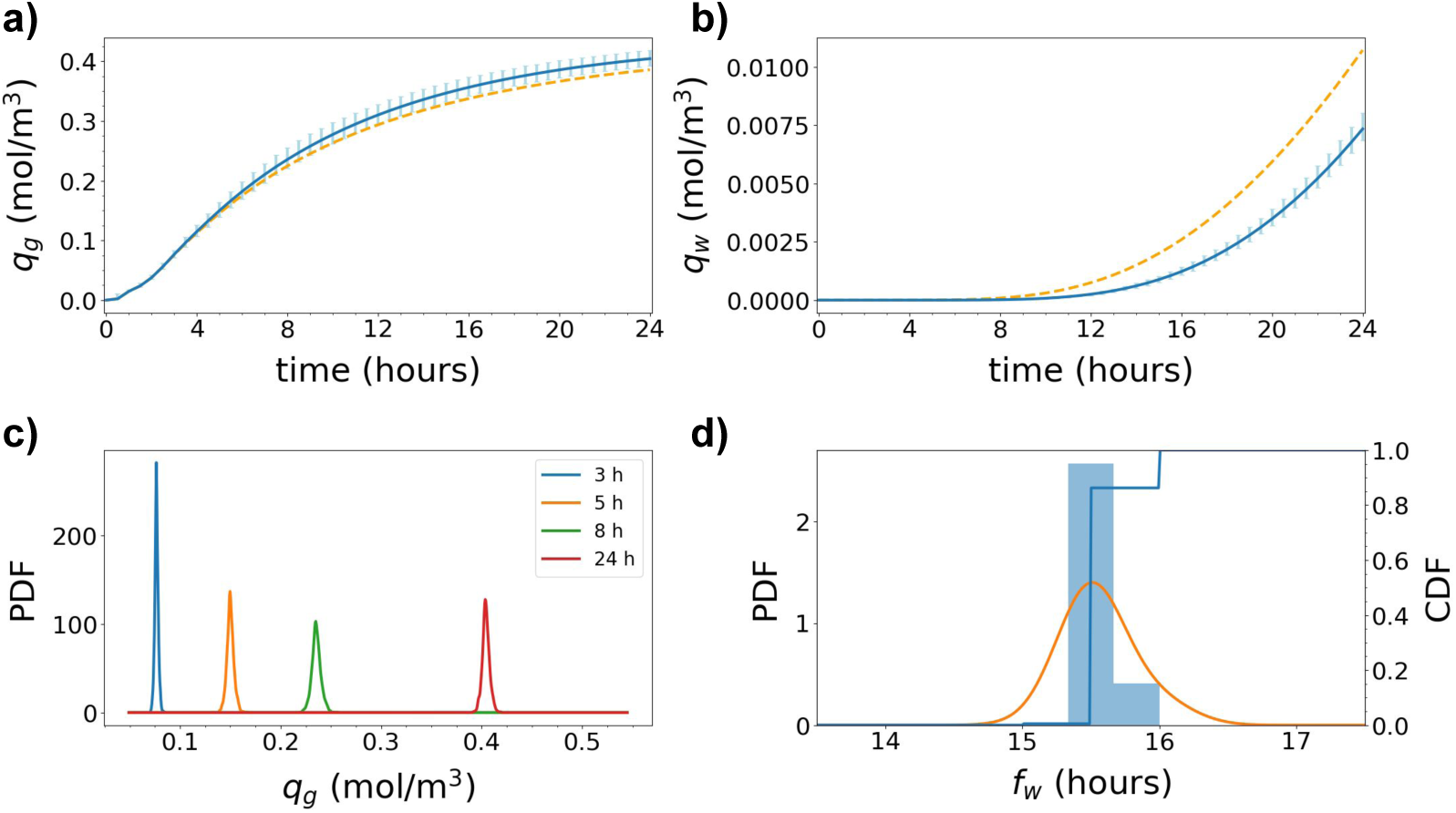
Uncertainty quantification for Model V1. The average tracer concentration in a subregion of (a) gray matter *q*_*g*_ and (b) white matter *q*_*w*_ as defined by (9). The blue curves show the expected value. The light blue vertical bars indicate the variability: 99.73% of the samples fall within the plotted range (with 0.135% of the samples above and 0.135% below). The dashed orange lines in (a) and (b) indicate the analogous expected value curve resulting from Model D1 (constant diffusion only), for comparison. Expected values for *q*_*g*_ are nearly identical as for Model D1 and D2, but variation is much lower. Expected values for *q*_*w*_ are lower than for Model D1 and variation is much lower (c) The probability density functions (PDFs) corresponding to *q*_*g*_ at 3, 5, 8 and 24 hours after tracer injection. The PDFs show very low variation. Variation increases slightly over time. (d) Histogram of white subregion activation time *f*_*w*_ as defined by (11) (bars), corresponding estimated PDF (orange curve), and corresponding cumulative density function (CDF).

**FIGURE 9.**
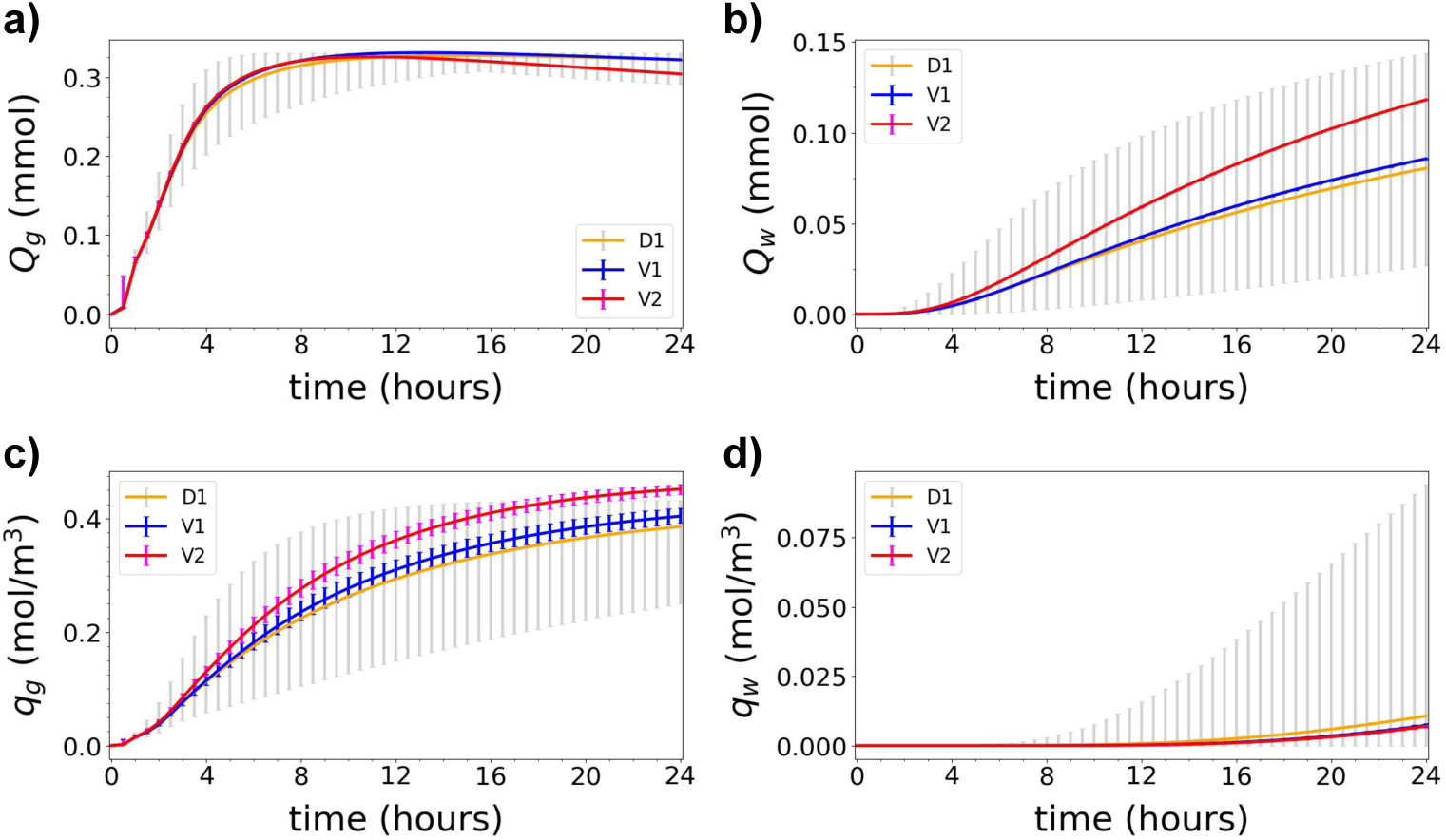
Uncertainty quantification for Model V2. Model V2 (red) in comparison with Models D1 (orange) and V1 (blue). The integrated amount of tracer in the (a) gray matter *Q*_*g*_ and (b) white matter *Q*_*w*_, as defined by (8), over time. The average tracer concentration in a subregion of (c) gray matter *q*_*g*_ and (d) white matter *q*_*w*_, as defined by (9), over time. The curves show the expected values while vertical bars indicate the 99.73% confidence intervals of the different models.

When considering the average concentration of tracer in two smaller regions of interest (cf. (9)), variability in model D2 increases drastically (Figure 7). In the gray matter region (Figure 7a), the expected average tracer concentration increases steadily to 0.11 mol/m^3^ after 4 hours, 0.23 mol/m^3^ after 8 hours, 0.35 mol/m^3^ after 16 hours and is still increasing after 24 hours. The variability is moderate after 3 hours (Figure 7c), but increases thereafter. The length of the 99.73% confidence interval peaks at 0.39 mol/m^3^ after 11 hours before decreasing moderately for later times.

The expected average tracer concentration in the white matter is low, lower than in the gray matter (Figure 7b) by a factor of at least 40, and starts increasing only after approximately 14 hours. For the samples in the lower range of the 99.73% interval (thus with the lower effective diffusivity), the concentration in the white matter region remains close to zero after 24 hours. For the white region activation time, we observe some variability (Figure 7d): the peak likelihood is after 14-15 hours, less (than 10%) likely to be less than 12 hours, and (more than 90%) likely to be less than 19 hours. The white subregion activation threshold was reached in all samples within the simulation time span.

### Quantifying the effect of glymphatic circulation

In light of the substantial uncertainty surrounding ISF/CSF flow in paravascular/perivascular spaces and potential ISF flow in extracellular spaces, we now turn to study the effect of uncertain velocity fields. To investigate the effect of uncertainty in a glymphatic velocity model, we defined a random velocity field with correlation length corresponding to the typical distance between parenchymal arterioles and venules (Model V1).

The expected amounts of tracer found in the whole gray and whole white matter for Model V1 are nearly identical to those found for Model D2 and Model D1, while the variability is minimal (data shown later cf. Figure 9). Thus, on average, small random variations in fluid velocity did not increase (or decrease) the tracer distribution into the parenchyma on a global scale. This observation can be interpreted in the light of the small correlation length of the velocity field compared to the size of the whole gray and white matter.

The expected average tracer concentration in the gray subregion *q*_*g*_ reaches 0.2 mol/m^3^ in 7 hours (Figure 8a). This is a considerable amount of time, given that the initial average SAS concentration is 3.57 mol/m^3^. The expected average tracer concentration in the white subregion *q*_*w*_ is lower, and only reaches 7.3 mmol/m^3^ in 24 hours (Figure 8b). We observe that the expected *q*_*g*_ increases marginally faster with the glymphatic velocity model than for pure diffusion: at 24 hours, *q*_*g*_ is 2.5% higher for V1 (0.40 mol/m^3^) than for D1 (0.39 mol/m^3^). On the other hand, the expected *q*_*w*_ increases faster with pure diffusion than with the glymphatic velocity model: at 24 hours, *q*_*w*_ is 34% lower for V1 (0.0073 mol/m^3^) than for D1 (0.011 mol/m^3^). The peak relative difference between pure diffusion and the upper limit of the 99.73% interval of model V1 is high after one hour, due to low tracer concentration overall. The next peak occurs after 8 hours where the relative difference is 13 % between the two.

However, the variation in both gray and white local average tracer concentration is small. For early time points (up to 3-4 hours), nearly no variation is evident in the average tracer concentration of the local regions (Figure 8a-c). The peak length of the 99.73% interval for *q*_*g*_ is 0.035 mol/m^3^ (at 9 hours), and the relative variability ranges from 6-19% in the 24 hour time span.

Moreover, the activation time *f*_*w*_ shows low variability: all simulations resulted in an activation time of 15.5-16 hours (Figure 8d). The substantially reduced variability for V1 compared to e.g. D2 combined with the comparable expected values yields much larger likely sample ranges for D2 than for V1.

### Quantifying the effect of glymphatic directionality

The cardiovascular pulse propagates along the larger arteries entering the brain from below before spreading outwards [41, 58]. To assess whether and how such a directionality in the glymphatic system affects parenchymal tracer distribution, we added a net flow field to the random velocity field representing the glymphatic circulation (Model V2).

With more fluid entering the brain from below, as illustrated by the streamlines of Figure 2c, the total parenchymal amount of tracer increases. For the expected amount of tracer in gray matter, however, Model V2 was in very good agreement with Models D1 and V1 (Figure 9a). After 13 hours, the amount of tracer found in the gray matter is higher for Model D1 than for Model V2. In Model V2, more of the tracer is found deeper in the gray matter and eventually moves to the white matter. We note that the uncertainty associated with the velocity fields barely affects the amount of tracer in the gray and white matter, as demonstrated by the nearly vanishing variation associated with *Q*_*g*_ and *Q*_*w*_ for Model V2 (and V1) (Figure 9a-b).

The expected amount of tracer in the white matter *Q*_*w*_ increases substantially by the introduction of the directional velocity field (Figure 9b). The expected value curve starts deviating from the other models after 4-5 hours, and the difference increases with time. At 24 hours, the expected amount of tracer found in the white matter *Q*_*w*_ is 50% larger for Model V1 (0.12 mmol) as for Model D1 (0.08 mmol). However, in view of the large variability associated with *Q*_*w*_ for Model D1 and the nearly vanishing variability associated with Model V2, the expected amount of white matter tracer for Model V2 falls well within the 99.73% confidence interval for Model D1.

The directional velocity field also induces an increase in the expected average tracer concentration in the gray subregion *q*_*g*_ (0.45 mol/m^3^ vs 0.40 for V1 and 0.39 mmol/m^3^ for D1 at 24 hours, Figure 9c). In contrast to for *Q*_*g*_ and *Q*_*w*_, this functional also displays some variability, with a peak variability (0.031 mol/m^3^ i.e. 10%) at 8-10 hours after injection. Notably, after 21-22 hours, the average tracer concentration in gray matter is larger than for pure diffusion (and for no net flow) also in terms of 99.73% confidence intervals. For *q*_*w*_, Model V1 and V2 are in close agreement, both with distinctly less variability than Model D1 (Figure 9d).

### Quantifying the effect of paraarterial influx with drainage

A number of open questions remain in the context of glymphatic and paravascular efflux routes. To further investigate potential pathways, we also considered a model representing paraarterial influx combined with parenchymal ISF drainage (Model V3).

Paraarterial inflow with drainage increases the amount of tracer found in the parenchyma for the early time points (Figure 10). After 4 hours, with the lowest velocities, the amount of tracer in the gray matter is equal to models with only diffusion (0.25 mmol). With higher velocities, however, the amount of tracer found in the gray matter increases by 32% to reach 0.33 mmol. After a peak at 6-8 hours, drainage and transport into white matter cause a decrease in the expected amount of tracer in the gray matter, while its variation stays more or less constant (0.11-0.12 mmol). The PDFs of the amount of tracer found in the gray matter thus have different characteristics than the two previous models, in particular the red curve (24 hours) shows lower amounts of tracer than at the two previous time points.

**FIGURE 10.**
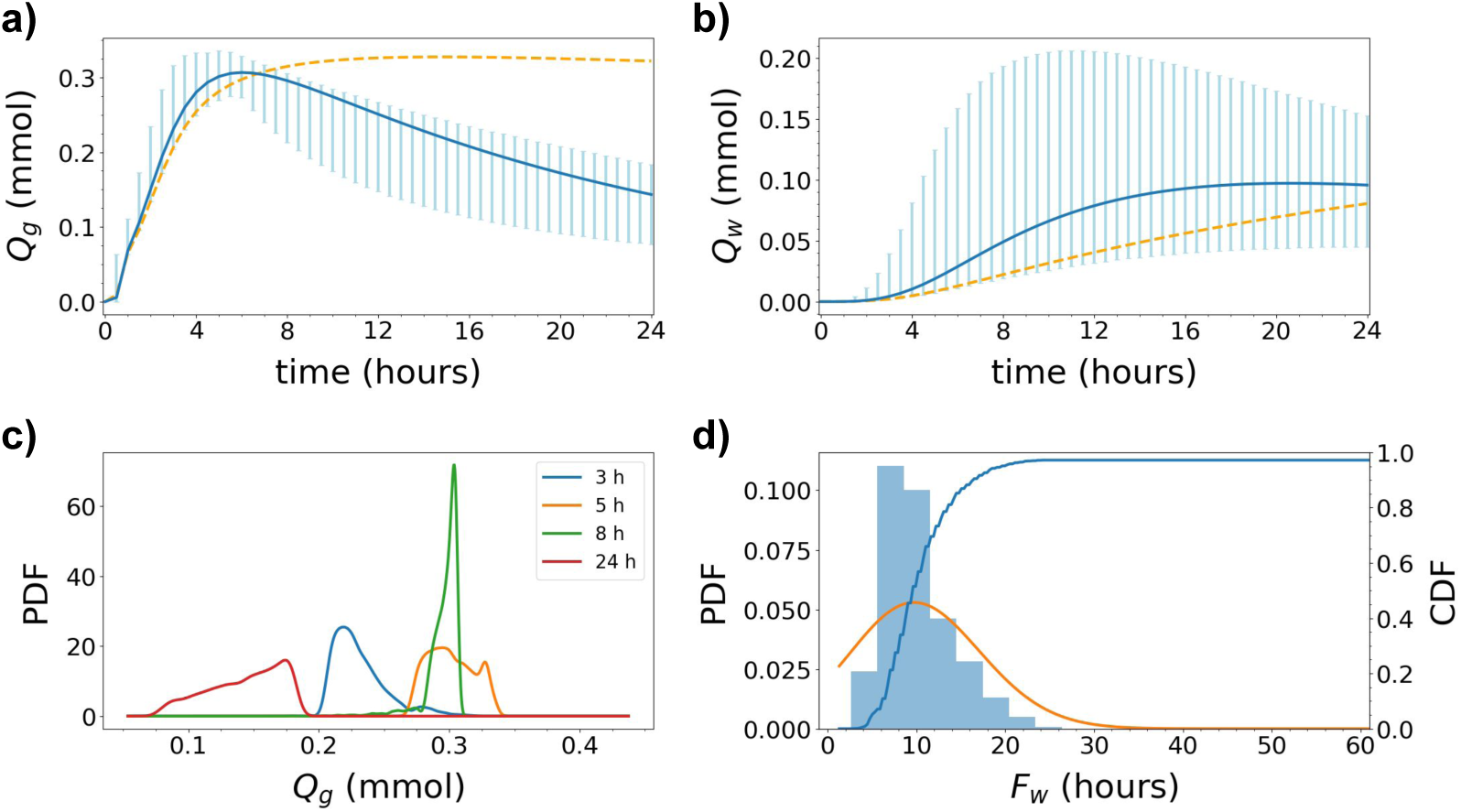
Uncertainty quantification for Model V3. The integrated amount of tracer in the (a) gray matter *Q*_*g*_ and (b) white matter *Q*_*w*_ over time; *Q*_*g*_ and *Q*_*w*_ as defined by (8). The blue curves show the expected value. The light blue vertical bars indicate the variability: 99.73% of the samples fall within the plotted range (with 0.135% of the samples above and 0.135% below). The dashed orange lines in (a) and (b) indicate the analogous expected value curve resulting from Model D1 (constant diffusion only), for comparison. Large variations in the white matter is found depending on the inflow velocity. (c) The probability density functions (PDFs) corresponding to *Q*_*g*_ at 3, 5, 8 and 24 hours after tracer injection. (d) Histogram of white matter activation time *F*_*w*_ as defined by (10) (bars), corresponding estimated PDF (orange curve), and corresponding cumulative density function (CDF). We note that the CDF peaks at 0.96 (¡ 1.0) as some samples never reached the white region activation threshold.

For the white matter, the expected amount of tracer increases with time, rapidly in comparison with pure diffusion, and seems to peak at approximately 0.097 mmol (at 19-22 hours) before slowly decreasing. Variation, on the other hand, is substantial and in some cases the amount of tracer found in the white matter reaches 0.2 mmol, which higher than what is seen in any previous model. This is visible by a peak of the maximum values within the 99.73% interval after 11-12 hours. In Model V3, tracer is drained out of the system and the amount of tracer in the white matter is similar as for the previous models at 24 hours.

The white matter activation time is likely lower for Model V3 compared to previous models, and the variation is substantial (Figure 10d). The white matter activation time is less (than 10%) likely to be less than 6 hours, but (more than 90%) likely to be less than 16.5 hours. Note that the white matter activation threshold was not reached in 3% of the samples.

## DISCUSSION

In this study, we have investigated the variability in parenchymal tracer enhancement resulting from uncertainty in diffusion and convection parameters. We designed five computational models representing different diffusion and convection regimes and used stochastic analysis to rigorously evaluate the resulting probability distributions.

In all models, 10% of the tracer reached the white matter within 40 hours, with more variability in activation time for diffusion models and less variability for models including a convective velocity. Indeed, uncertainty in the diffusion parameters had a substantial impact on the amount of tracer in gray and white matter, and on the average tracer concentration in gray and white subregions. Overall, diffusion was not sufficient, with high likelihood, to transport tracer deep into the parenchyma.

A stochastic velocity field representing the glymphatic theory did not increase transport into any of the regions considered, unless augmented with an additional net flow with a prescribed directionality. In the latter case, transport was increased with overwhelming likelihood: for model V2, the entire 99.73% confidence interval for the gray subregion average tracer concentration was higher than for model D1. Models including parenchymal drainage displayed substantial variability, and reached peak values for the expected amount of tracer both in gray and white matter within 24 hours.

### Comparison with previous work

Our models mimic the experimental set-up of an MRI study of parenchymal tracer distribution after intrathecal gadobutrol injection [64]. In our simulations, as in the MRI study, the tracer first spreads to inferior regions of the parenchyma closer to the (modelled) injection site. Modelling a healthy patient, we assumed that the tracer concentration in the ventricular CSF was low [64, 63]. Thus, no tracer spreads to the parenchyma from the ventricles directly. In models with diffusion only, the amount of tracer in the gray matter peaks at approximately 15 hours. In the MRI study, the time to peak enhancement in selected regions of interest was between 12 and 24 hours [64]. In a more recent study, time to peak values were considerably longer, up to 48 hours, for some regions [63]. However, in the latter study, the time to peak enhancement was shorter for the white matter than for the gray matter in healthy subjects. This observation is not consistent with the results from either of our computational models.

Most of the reported time to peak values in the two human MRI-studies [64, 63] are within the 99.73% confidence interval of the random homogeneous diffusion model (Model D1). However, even for the upper range of the confidence interval, the time to peak/steady state value for the white matter exceeds 24 hours in our model. The uncertainty in the diffusion coefficient may explain a four-fold difference in the amount of tracer found in the white matter at 24 hours. Despite this large variation, the discrepancy between simulations and experiments in white matter could not be explained by uncertainty in the diffusion parameter. This may suggest other mechanisms in addition to diffusion for tracer transport into deeper regions of the brain. According to paraarterial influx theories in general and the glymphatic theory in particular, tracer flows rapidly along and into the parenchymal PVS [37] distributing tracer to the gray matter. Hence, one may expect diffusion models to underestimate the amount of tracer in gray matter at a given time. However, is worth noting that we do not observe such an underestimation in our diffusion model, when compared to the experimental values [64]. In contrast, we do observe a delayed distribution of tracer in white matter.

Brain tissue is known to be both anisotropic and heterogeneous [51, 62, 77]. We found the variation due to spatial heterogeneity in the diffusion coefficient to be low. As the correlation length was small compared to the size of the the gray and white matter, a lack to tracer concentration in one local region was balanced by enhancement in a different local region. In addition, we note that representing the diffusion coefficient as a random variable or a random field yields the same expected value. Tracer distribution to large brain regions can thus be well approximated using an average diffusion constant if the spatial heterogeneity is present on a shorter length scale.

In models with convection, given a homogenized velocity of average magnitude 0.17 *µ*m/s, tracer distribution depends on the characteristics of the velocity field. In the glymphatic theory, CSF enters the brain along arteries and re-enters the SAS along a paravenous efflux pathway [39, 37]. In our glymphatic circulation model, the stochastic velocity field, representing homogenized paraarterial and paravenous flow, did not increase tracer distribution to the brain. An increase in the amount of tracer surrounding paraarterial spaces was balanced by a lower distribution around paravenous spaces. However, when local regions are addressed, tracer concentration may increase by up to 13% compared to diffusion alone, depending on the surrounding velocity field and region of interest. As we consider a homogenized representation of the PVS, this change reflects an increase in regions surrounding arterial PVS (not only inside the PVS). Iliff et al. [38] reported a 2-fold increase in tracer intensity in PVS in normal mice compared to mice with internal carotid artery ligation. The increase in the surrounding parenchyma was lower, approximately 30-40%, which compares more naturally with our estimate of 13 %. It should be noted however, that our region of interest was deeper into the parenchyma (extending from 0.6 to 4 mm depth) than the region of interest (at 100 *µ*m) used by Iliff et al. [38]. Moreover, our model parameters reflect a different species (man versus mouse), and the tracer spread takes place at a longer time scale.

When modelling paraarterial influx combined with parenchymal drainage (Model V3), the time to peak was reduced to 6-8 hours in the gray matter. Although lacking quantitative drainage parameters, we observe that substantial clearance would reduce both the time to peak and relative tracer enhancement in the brain compared to diffusion alone. In the glymphatic directionality model (Model V2), guided by [58], the presence of a paravascular directional velocity also decreases the expected time to peak tracer enhancement in gray matter, down to 11 hours (compared to 15 hours for pure diffusion). Thus, when experimental data suggests a time to peak enhancement shorter than for diffusion alone, it is not clear whether this is due to increased glymphatic function or increased clearance by parenchymal drainage.

In our models, the white matter (and subregions) is where the effect of a convective velocity becomes most prominent. The only model modification causing an expected time to peak enhancement in white matter of approximately 24 hours is with a paraarterial inflow and drainage (Model V3). In this model, the upper limit of the 99.73% confidence interval peaks at approximately 12 hours, which is more comparable to the rapid tracer enhancement observed in the white matter of healthy subjects [63].

Although diffusion may act as the main transport mechanism in the parenchyma [68, 36], we here show that convective velocities of magnitude less than 1 *µ* m/s may play an important role for transport. This result holds when there is a structure of the glymphatic circulation as used in Model V2 or possibly a net inflow as in Model V3. It should be noted that this directional velocity field, in which pulsations propagate upwards from the brain stem [41, 58], favors inflow when tracer is injected in lower CSF regions such as e.g. in the spinal canal.

### Limitations

In the present study, we have used a continuous model of the brain parenchyma allowing only for an homogenized representation of paravascular spaces on the scale of micrometers. To remedy this limitation, combined with restrictions placed by mesh resolution, we used lower velocities acting over larger areas to model paravascular flows.

Further, we did not distinguish between white and gray matter in terms of the fluid velocity or in the diffusivity, although white matter is assumed to be more permeable [56]. However, in the absence of substantial drainage, net movement of fluid (in gray matter and PVS vs white matter) should on average be equal in the two regions by conservation of mass. Therefore, we used maximal velocity magnitudes of approximately 0.5 *µ*m/s, which is similar to what has been reported in white matter [1], but not as high as has been reported in local regions in the PVS [14, 48]. While we used qualitative measurements [41, 58] to suggest a directionality in the glymphatic circulation, we predict that more detailed measurements of glymphatic function in different brain regions would be important for tracer enhancement and clearance.

The boundary concentration in our model was assumed to spread in a manner similar to what was seen from the signal intensity in the MRI study by Ringstad et al. [64]. A more detailed analysis of the spread of tracer in the CSF could be based on at least solving the Navier-Stokes equations in the SAS. In addition, our model ignores other efflux pathways directly from the SAS such as e.g. arachnoid granulations [29], dural lymphatics [46, 45], and nasal lymphatics [49], although lymphatic drainage of CSF has recently been proposed to dominate glymphatic clearance [47].

In the experiments by Ringstad et al. [64, 63], tracer distribution within the parenchyma varied considerably from patient to patient. In our analysis, we did not consider patient-specific meshes, but rather one representative mesh. Patient-specific meshes would add additional dimensions to the space of uncertainty, possibly giving different distributions in output in each of the patients.

The MRI-studies [64, 63] only provide quantitative values of tracer enhancement signal intensity, and not tracer concentrations. As the relation between signal intensity and concentration is nonlinear [20], we have not made a direct comparison between these two quantities. However, we have assumed that a peak in signal intensity corresponds to a peak in tracer concentration, thus allowing for a comparison of time-to-peak between the model results and experiments.

## CONCLUSIONS

The results from this study show that uncertainty in the diffusion parameters substantially impact the amount of tracer in gray and white matter, and the average tracer concentration in gray and white subregions. However, even with an uncertainty in the diffusion coefficient of a factor three, and a resulting four-fold variation in white matter tracer enhancement, discrepancies between simulations of diffusion and experimental data are too large to be attributed to uncertainties in the diffusion coefficient alone.

A convective velocity field, representing the glymphatic circulation, increases tracer enhancement in the brain as compared to pure diffusion. However, increased enhancement is reliant on a directional structure of the velocity field allowing for efficient inflow from given regions.

Diffusion alone was able to mimic behaviour in MR-studies in specific regions. However, this result does not imply lack of glymphatic circulation as the gray matter tracer enhancement was equal for the glymphatic model with directionality and for diffusion alone. However, the white matter concentration was greatly increased in the former model. Thus measuring glymphatic function requires detailed experimental data and analysis of the whole brain.

## LIST OF ABBREVIATIONS

CDF: Cumulative density function
CSF: Cerebrospinal fluid
ISF: Interstitial fluid
MR(I): Magnetic resonance (imaging)
MC: Monte Carlo
PDE: Partial differential equation
PDF: Probability density function.
PVS: Paravascular/perivascular space(s)
SAS: Subarachnoid space
UQ: Uncertainty quantification

## DECLARATIONS

### Availability of data and material

The datasets generated and analyzed during the current study are available via the Uncertainty quantification of parenchymal tracer distribution using random diffusion and convective velocity fields (data sets): https://doi.org/10.5281/zenodo. 3241364. Additional data and computer code are available from the corresponding author on reasonable request.

### Authors’ contributions

VV, MC and MER designed the study. VV and MC conducted the experiments and analyzed the results. VV, MC and MER created the figures. VV, MC and MER wrote and reviewed the manuscript.

## Acknowledgements

The authors thank Patrick E. Farrell and Michael B. Giles (University of Oxford) for constructive discussions on topics related to the manuscript.

## References

[1] N. J. Abbott. Evidence for bulk flow of brain interstitial fluid: significance for physiology and pathology. Neurochemistry international, 45(4):545–552, 2004.

[2] N. J. Abbott, M. E. Pizzo, J. E. Preston, D. Janigro, and R. G. Thorne. The role of brain barriers in fluid movement in the CNS: is there a ‘glymphatic’ system? Acta neuropathologica, pages 1–21, 2018.

[3] P. Abrahamsen. A Review of Gaussian Random Fields and Correlation Functions. Norwegian Computing Center, 2 edition, 1997.

[4] D. L. Adams, V. Piserchia, J. R. Economides, and J. C. Horton. Vascular supply of the cerebral cortex is specialized for cell layers but not columns. Cerebral Cortex, 25(10):3673–3681, 2015.

[5] N. J. Albargothy, D. A. Johnston, M. MacGregor-Sharp, R. O. Weller, A. Verma, C. A. Hawkes, and R. O. Carare. Convective influx/glymphatic system: tracers injected into the csf enter and leave the brain along separate periarterial basement membrane pathways. Acta neuropathologica, 136(1):139–152, 2018.

[6] R. Aldea, R. O. Weller, D. M. Wilcock, R. O. Carare, and G. Richardson. Cerebrovascular smooth muscle cells as the drivers of intramural periarterial drainage of the brain. Frontiers in aging neuroscience, 11, 2019.

[7] M. Alnæs, J. Blechta, J. Hake, A. Johansson, B. Kehlet, A. Logg, C. Richardson, J. Ring, M. E. Rognes, and G. N. Wells. The FEniCS project version 1.5. Archive of Numerical Software, 3(100):9–23, 2015.

[8] M. Asgari, D. D. Zélicourt, and V. Kurtcuoglu. Glymphatic solute transport does not require bulk flow. Scientific reports, 6:38635, 2016.

[9] A. Aspelund, S. Antila, S. T. Proulx, T. V. Karlsen, S. Karaman, M. Detmar, H. Wiig, and K. Alitalo. A dural lymphatic vascular system that drains brain interstitial fluid and macromolecules. Journal of Experimental Medicine, 212(7):991–999, 2015.

[10] R. N. Auer. Histopathology of brain tissue response to stroke and injury. In Stroke (Sixth Edition), pages 47–59. Elsevier, 2016.

[11] E. N. Bakker, D. M. Naessens, and E. VanBavel. Paravascular spaces: entry to or exit from the brain? Experimental physiology, 2018.

[12] S. Balay, S. Abhyankar, M. Adams, J. Brown, P. R. Brune, K. Buschelman, V. Eijkhout, W. Gropp, D. Kaushik, M. G. Knepley, and Others. PETSc users manual revision 3.8. Technical report, Argonne National Laboratory, ANL, 2017.

[13] O. Balédent, C. Gondry-Jouet, M.-E. Meyer, G. De Marco, D. Le Gars, M.-C. Henry-Feugeas, and I. Idy-Peretti. Relationship between cerebrospinal fluid and blood dynamics in healthy volunteers and patients with communicating hydrocephalus. Investigative radiology, 39(1):45–55, 2004.

[14] B. Bedussi, M. Almasian, J. de Vos, E. VanBavel, and E. N. Bakker. Paravascular spaces at the brain surface: Low resistance pathways for cerebrospinal fluid flow. Journal of Cerebral Blood Flow & Metabolism, 38(4):719–726, 2017.

[15] B. Bedussi, M. G. Lier, J. W. Bartstra, J. Vos, M. Siebes, E. VanBavel, and E. N. Bakker. Clearance from the mouse brain by convection of interstitial fluid towards the ventricular system. Fluids and Barriers of the CNS, 12(1):23, 2015.

[16] B. Bedussi, N. N. van der Wel, J. de Vos, H. van Veen, M. Siebes, E. VanBavel, and E. N. Bakker. Paravascular channels, cisterns, and the subarachnoid space in the rat brain: A single compartment with preferential pathways. Journal of Cerebral Blood Flow & Metabolism, 37(4):1374–1385, 2017.

[17] J. Biehler, M. W. Gee, and W. A. Wall. Towards efficient uncertainty quantification in complex and large-scale biomechanical problems based on a Bayesian multi-fidelity scheme. Biomech Model Mechanobiol, 14:489–513, 2015.

[18] R. Carare, M. Bernardes-Silva, T. Newman, A. Page, J. Nicoll, V. Perry, and R. Weller. Solutes, but not cells, drain from the brain parenchyma along basement membranes of capillaries and arteries: significance for cerebral amyloid angiopathy and neuroimmunology. Neuropathology and applied neurobiology, 34(2):131–144, 2008.

[19] J. Charrier, R. Scheichl, and A. L. Teckentrup. Finite element error analysis of elliptic PDEs with random coefficients and its application to multilevel Monte Carlo methods. SIAM Journal of Numerical Analysis, 51(1):322–352, 2013.

[20] X. Chen, G. W. Astary, H. Sepulveda, T. H. Mareci, and M. Sarntinoranont. Quantitative assessment of macromolecular concentration during direct infusion into an agarose hydrogel phantom using contrast-enhanced mri. Magnetic resonance imaging, 26(10):1433–1441, 2008.

[21] K. A. Cliffe, M. B. Giles, R. Scheichl, and A. L. Teckentrup. Multilevel Monte Carlo methods and applications to elliptic PDEs with random coefficients. Computing and Visualization in Science, 14(1):3–15, 2011.

[22] M. Croci, M. B. Giles, M. E. Rognes, and P. E. Farrell. Efficient White Noise Sampling and Coupling for Multilevel Monte Carlo with Nonnested Meshes. SIAM/ASA Journal on Uncertainty Quantification, 6(4):1630–1655, 2018.

[23] A. K. Diem, M. MacGregor Sharp, M. Gatherer, N. W. Bressloff, R. O. Carare, and G. Richardson. Arterial pulsations cannot drive intramural periarterial drainage: significance for a*β* drainage. Frontiers in neuroscience, 11:475, 2017.

[24] H. C. Elman, D. J. Silvester, and A. J. Wathen. Finite elements and fast iterative solvers: with applications in incompressible fluid dynamics. Oxford University Press, USA, 2014.

[25] R. D. Falgout and U. M. Yang. Hypre: A library of high performance preconditioners. In International Conference on Computational Science, pages 632–641. Springer, Springer, 2002.

[26] Q. Fang. Mesh-based monte carlo method using fast ray-tracing in plücker coordinates. Biomedical optics express, 1(1):165–175, 2010.

[27] C. Geuzaine and J.-F. Remacle. Gmsh: A 3-d finite element mesh generator with built-in pre- and postprocessing facilities. International journal for numerical methods in engineering, 79(11):1309–1331, 2009.

[28] L. Guo, J. C. Vardakis, T. Lassila, M. Mitolo, N. Ravikumar, D. Chou, M. Lange, A. Sarrami-Foroushani,B. J. Tully, Z. A. Taylor, et al. Subject-specific multi-poroelastic model for exploring the risk factors associated with the early stages of alzheimer’s disease. Interface focus, 8(1):20170019, 2018.

[29] A. C. Guyton and J. E. Hall. Textbook of medical physiology. 11th. WB Sounders Company, Philadelphia, 2006.

[30] P. Hadaczek, Y. Yamashita, H. Mirek, L. Tamas, M. C. Bohn, C. Noble, J. W. Park, and K. Bankiewicz. The “perivascular pump” driven by arterial pulsation is a powerful mechanism for the distribution of therapeutic molecules within the brain. Molecular Therapy, 14(1):69–78, 2006.

[31] M.-J. Hannocks, M. E. Pizzo, J. Huppert, T. Deshpande, N. J. Abbott, R. G. Thorne, and L. Sorokin. Molecular characterization of perivascular drainage pathways in the murine brain. Journal of Cerebral Blood Flow & Metabolism, 38(4):669–686, 2018.

[32] I. F. Harrison, B. Siow, A. B. Akilo, P. G. Evans, O. Ismail, Y. Ohene, P. Nahavandi, D. L. Thomas, M. F. Lythgoe, and J. A. Wells. Non-invasive imaging of csf-mediated brain clearance pathways via assessment of perivascular fluid movement with dti mri. eLife, 7:e34028, 2018.

[33] P. Hauseux, J. S. Hale, S. Cotin, and S. P. A. Bordas. Quantifying the uncertainty in a hyperelastic soft tissue model with stochastic parameters. Applied Mathematical Modelling, 62:86–102, oct 2018.

[34] S. B. Hladky and M. A. Barrand. Mechanisms of fluid movement into, through and out of the brain: evaluation of the evidence. Fluids and Barriers of the CNS, 11(1):26, 2014.

[35] S. B. Hladky and M. A. Barrand. Elimination of substances from the brain parenchyma: efflux via perivascular pathways and via the blood–brain barrier. Fluids and Barriers of the CNS, 15(1):30, 2018.

[36] K. E. Holter, B. Kehlet, A. Devor, T. J. Sejnowski, A. M. Dale, S. W. Omholt, O. P. Ottersen, E. A. Nagelhus, K.-A. Mardal, and K. H. Pettersen. Interstitial solute transport in 3D reconstructed neuropil occurs by diffusion rather than bulk flow. Proceedings of the National Academy of Sciences, 114(37):9894–9899, 2017.

[37] J. J. Iliff, M. Wang, Y. Liao, B. A. Plogg, W. Peng, G. A. Gundersen, et al. A paravascular pathway facilitates CSF flow through the brain parenchyma and the clearance of interstitial solutes, including amyloid beta. Sci Transl Med, 4(147):147ra111, 2012.

[38] J. J. Iliff, M. Wang, D. M. Zeppenfeld, A. Venkataraman, B. A. Plog, Y. Liao, R. Deane, and M. Nedergaard. Cerebral arterial pulsation drives paravascular csf–interstitial fluid exchange in the murine brain. Journal of Neuroscience, 33(46):18190–18199, 2013.

[39] N. A. Jessen, A. S. F. Munk, I. Lundgaard, and M. Nedergaard. The glymphatic system: a beginner’s guide. Neurochemical research, 40(12):2583–2599, 2015.

[40] B.-J. Jin, A. J. Smith, and A. S. Verkman. Spatial model of convective solute transport in brain extracellular space does not support a ‘glymphatic’ mechanism. The Journal of general physiology, 148(6):489–501, 2016.

[41] V. Kiviniemi, X. Wang, V. Korhonen, T. Keinänen, T. Tuovinen, J. Autio, P. LeVan, S. Keilholz, Y.-F. Zang, J. Hennig, et al. Ultra-fast magnetic resonance encephalography of physiological brain activity–glymphatic pulsation mechanisms? Journal of Cerebral Blood Flow & Metabolism, 36(6):1033–1045, 2016.

[42] J. Krejza, M. Arkuszewski, S. E. Kasner, J. Weigele, A. Ustymowicz, R. W. Hurst, B. L. Cucchiara, and S. R. Messe. Carotid artery diameter in men and women and the relation to body and neck size. Stroke, 37(4):1103–1105, 2006.

[43] P. Lacolley, P. Challande, S. Boumaza, G. Cohuet, S. Laurent, P. Boutouyrie, J.-A. Grimaud, D. Paulin, J.-M. D. Lamaziere, and Z. Li. Mechanical properties and structure of carotid arteries in mice lacking desmin. Cardiovascular research, 51(1):178–187, 2001.

[44] F. Lindgren, H. Rue, and J. Lindström. An explicit link between Gaussian fields and Gaussian Markov random fields: the stochastic partial differential equation approach. Journal of the Royal Statistical Society: Series B (Statistical Methodology), 73(4):423–498, 2009.

[45] A. Louveau, B. A. Plog, S. Antila, K. Alitalo, M. Nedergaard, and J. Kipnis. Understanding the functions and relationships of the glymphatic system and meningeal lymphatics. The Journal of clinical investigation, 127(9):3210–3219, 2017.

[46] A. Louveau, I. Smirnov, T. J. Keyes, J. D. Eccles, S. J. Rouhani, J. D. Peske, N. C. Derecki, D. Castle, J. W. Mandell, K. S. Lee, et al. Structural and functional features of central nervous system lymphatic vessels. Nature, 523(7560):337, 2015.

[47] Q. Ma, M. Ries, Y. Decker, A. Müller, C. Riner, A. Bücker, K. Fassbender, M. Detmar, and S. T. Proulx. Rapid lymphatic efflux limits cerebrospinal fluid flow to the brain. Acta neuropathologica, pages 1–15, 2019.

[48] H. Mestre, J. Tithof, T. Du, W. Song, W. Peng, A. M. Sweeney, G. Olveda, J. H. Thomas, M. Nedergaard, and D. H. Kelley. Flow of cerebrospinal fluid is driven by arterial pulsations and is reduced in hypertension. Nature communications, 9(1):4878, 2018.

[49] R. Mollanji, R. Bozanovic-Sosic, A. Zakharov, L. Makarian, and M. G. Johnston. Blocking cerebrospinal fluid absorption through the cribriform plate increases resting intracranial pressure. American Journal of Physiology-Regulatory, Integrative and Comparative Physiology, 2002.

[50] R. B. Nelsen. An introduction to copulas. Springer Science & Business Media, 2007.

[51] C. Nicholson. Diffusion and related transport mechanisms in brain tissue. Reports on progress in Physics, 64(7):815, 2001.

[52] C. Nilsson, F. Stahlberg, C. Thomsen, O. Henriksen, M. Herning, and C. Owman. Circadian variation in human cerebrospinal fluid production measured by magnetic resonance imaging. American Journal of Physiology-Regulatory, Integrative and Comparative Physiology, 262(1):R20–R24, 1992.

[53] D. Orešković, M. Radoš, and M. Klarica. New concepts of cerebrospinal fluid physiology and development of hydrocephalus. Pediatric neurosurgery, 52(6):417–425, 2017.

[54] D. Orešković, M. Radoš, and M. Klarica. Role of choroid plexus in cerebrospinal fluid hydrodynamics. Neuroscience, 354:69–87, 2017.

[55] R. Potsepaev and C. L. Farmer. Application of stochastic partial differential equations to reservoir property modelling. In ECMOR XII-12th European Conference on the Mathematics of Oil Recovery, volume 2, 2014.

[56] S. S. Prabhu, W. C. Broaddus, G. T. Gillies, W. G. Loudon, Z.-J. Chen, and B. Smith. Distribution of macro-molecular dyes in brain using positive pressure infusion: a model for direct controlled delivery of therapeutic agents. Surgical neurology, 50(4):367–375, 1998.

[57] A. Quaglino, S. Pezzuto, and R. Krause. Generalized Multifidelity Monte Carlo Estimators. Preprint, 2018.

[58] Z. Rajna, L. Raitamaa, T. Tuovinen, J. Heikkilä, V. Kiviniemi, and T. Seppänen. 3d multi-resolution optical flow analysis of cardiovascular pulse propagation in human brain. IEEE transactions on medical imaging, 2019.

[59] L. Ray, J. J. Iliff, and J. J. Heys. Analysis of convective and diffusive transport in the brain interstitium. Fluids and Barriers of the CNS, 16(1):6, 2019.

[60] M. Rennels, O. Blaumanis, and P. Grady. Rapid solute transport throughout the brain via paravascular fluid pathways. Advances in neurology, 52:431–439, 1990.

[61] M. L. Rennels, T. F. Gregory, O. R. Blaumanis, K. Fujimoto, and P. A. Grady. Evidence for a ‘paravascular’fluid circulation in the mammalian central nervous system, provided by the rapid distribution of tracer protein throughout the brain from the subarachnoid space. Brain research, 326(1):47–63, 1985.

[62] M. E. Rice, Y. C. Okada, and C. Nicholson. Anisotropic and heterogeneous diffusion in the turtle cerebellum: implications for volume transmission. Journal of neurophysiology, 70(5):2035–2044, 1993.

[63] G. Ringstad, L. M. Valnes, A. M. Dale, A. H. Pripp, S.-A. S. Vatnehol, K. E. Emblem, K.-A. Mardal, and P. K. Eide. Brain-wide glymphatic enhancement and clearance in humans assessed with mri. JCI insight, 3(13), 2018.

[64] G. Ringstad, S. A. S. Vatnehol, and P. K. Eide. Glymphatic mri in idiopathic normal pressure hydrocephalus. Brain, 140(10):2691–2705, 2017.

[65] G. Roth and U. Dicke. Evolution of the brain and intelligence. Trends in cognitive sciences, 9(5):250–257, 2005.

[66] M. K. Sharp, R. O. Carare, and B. A. Martin. Dispersion in porous media in oscillatory flow between flat plates: applications to intrathecal, periarterial and paraarterial solute transport in the central nervous system. Fluids and Barriers of the CNS, 16(1):13, 2019.

[67] A. J. Smith and A. S. Verkman. The ‘glymphatic’ mechanism for solute clearance in alzheimer’s disease: game changer or unproven speculation? The FASEB Journal, 32(2):543–551, 2017.

[68] A. J. Smith, X. Yao, J. A. Dix, B.-J. Jin, and A. S. Verkman. Test of the ‘glymphatic’ hypothesis demonstrates diffusive and aquaporin-4-independent solute transport in rodent brain parenchyma. Elife, 6:e27679, 2017.

[69] J. M. Tarasoff-Conway, R. O. Carare, R. S. Osorio, L. Glodzik, T. Butler, E. Fieremans, L. Axel, H. Rusinek, C. Nicholson, B. V. Zlokovic, et al. Clearance systems in the brain – implications for alzheimer disease. Nature reviews neurology, 11(8):457, 2015.

[70] A. L. Teckentrup, R. Scheichl, M. B. Giles, and E. Ullmann. Further analysis of multilevel Monte Carlo methods for elliptic PDEs with random coefficients. Numerische Mathematik, 125:569–600, 2013.

[71] V. Thomée. On positivity preservation in some finite element methods for the heat equation. In International Conference on Numerical Methods and Applications, pages 13–24. Springer, 2014.

[72] D. S. Tuch, T. G. Reese, M. R. Wiegell, N. Makris, J. W. Belliveau, and V. J. Wedeen. High angular resolution diffusion imaging reveals intravoxel white matter fiber heterogeneity. Magnetic Resonance in Medicine: An Official Journal of the International Society for Magnetic Resonance in Medicine, 48(4):577–582, 2002.

[73] G. F. Vindedal, A. E. Thoren, V. Jensen, A. Klungland, Y. Zhang, M. J. Holtzman, O. P. Ottersen, and E. A. Nagelhus. Removal of aquaporin-4 from glial and ependymal membranes causes brain water accumulation. Molecular and Cellular Neuroscience, 77:47–52, 2016.

[74] J. H. Wood. Neurobiology of cerebrospinal fluid 2. Springer Science & Business Media, 2013.

[75] L. Xie, H. Kang, Q. Xu, M. J. Chen, Y. Liao, M. Thiyagarajan, J. O’Donnell, D. J. Christensen, C. Nicholson, J. J. Iliff, et al. Sleep drives metabolite clearance from the adult brain. science, 342(6156):373–377, 2013.

[76] Z. Ye, Y. Liu, X. Wang, X. Chen, C. Lin, Y. Su, T. Wang, Y. Tang, Y. Wu, and C. Qin. A rhesus monkey model of common carotid stenosis. Int J Clin Exp Med, 9(9):17487–17497, 2016.

[77] S. Zuzana, E. Syková, et al. Diffusion heterogeneity and anisotropy in rat hippocampus. Neuroreport, 9(7):1299–1304, 1998.

